# Small habitat patches can be the largest contributors to urban biodiversity across taxonomic groups

**DOI:** 10.64898/2026.01.26.701683

**Authors:** Kilian Perrelet, Lauren M. Cook, Florian Altermatt, Federico Riva, Marco Moretti

**Affiliations:** Swiss Federal Research Institute for Forest, Snow, and Landscape Research (WSL), Biodiversity and Conservation Biology, Birmensdorf, Switzerland; Swiss Federal Institute of Aquatic Science & Technology (Eawag), Department of Urban Water Management, Dübendorf, Switzerland; University of Zurich, Department of Evolutionary Biology and Environmental Studies, Zurich, Switzerland; Swiss Federal Institute of Aquatic Science & Technology (Eawag), Department of Aquatic Ecology, Dübendorf, Switzerland; Vrije Universiteit Amsterdam, Institute of Environmental Studies, Amsterdam, the Netherlands

**Keywords:** Invertebrates, Vertebrates, Plants, SLOSS, Urban green spaces, Patch size

## Abstract

**Aim:** As cities densify and expand, the careful planning and design of green spaces are essential for supporting urban biodiversity. Here, we evaluate the relative contribution of habitat patches of varying size, quality, and connectivity to urban biodiversity and assess environmental factors driving differences in species richness and community composition.

**Location:** Zurich, Switzerland.

**Time period: 2008–2018.:** Major taxa studied: Invertebrates, vertebrates, and trees.

**Methods:** We analyzed species occupancy data from 452 habitat patches. We quantified alpha, beta, and gamma diversity, assessed species-area relationships, and applied generalized dissimilarity modelling to test the role of patch area, connectivity, and habitat quality—proxied through environmental variables, including vegetation complexity, water presence, and forest isolation—in shaping community composition.

**Results:** Alpha diversity increased significantly with area, although small patches (usually < 5 ha) disproportionately contributed to beta diversity. Per unit area, groups of small patches yielded higher gamma diversity than equivalent areas of large patches, particularly for trees and invertebrates. Community composition was strongly influenced by patch area, with effects mediated by vegetation complexity, water, and isolation, with responses differing among taxa.

**Main conclusions:** Small habitat patches play a critical role in enhancing overall urban biodiversity. They increase species richness through cumulative area effects and promote community turnover (mediated by environmental heterogeneity). Maintaining networks of small patches alongside large green spaces is therefore key to conserving biodiversity in urban landscapes.

## Introduction

Urban areas are expected to accommodate up to 70% of the world’s population by 2050 and are expanding rapidly worldwide (United Nations, 2018). Among the most important ecological consequences of this expansion are habitat loss and fragmentation, which, when co-occurring, are typically linked to biodiversity decline (Fahrig, 2017; Keck et al., 2025; Simkin et al., 2022). While habitat loss consistently reduces biodiversity, the ecological consequences of habitat fragmentation *per se*—differences in habitat area and configuration independent of total habitat amount—remain debated (Fahrig et al., 2022; Fletcher et al., 2018; Riva, Haddad, et al., 2024; Riva, Koper, et al., 2024). In urban ecosystems, where both habitat loss and fragmentation are often extreme, understanding how patches of different sizes contribute to biodiversity is essential for guiding the design and spatial planning of both existing and novel habitats created through human activity (e.g., blue-green infrastructure) (Alberti et al., 2020; Aronson et al., 2016; Perrelet et al., 2024).

A central question in conservation and management of landscapes is whether one large or several smaller patches of equal total area support greater biodiversity—a long-standing topic known as the Single Large Or Several Small (SLOSS) question (Diamond, 1975; Fahrig, 2020; Fahrig et al., 2022; Simberloff & Abele, 1976). While species-area relationships (SARs) predict higher local (alpha) diversity in larger patches due to greater habitat and resource availability (MacArthur & Wilson, 1967), landscape-scale (gamma) biodiversity does not depend solely on patch-scale diversity patterns (Perrin et al., 2025; Riva & Fahrig, 2023a). In fact, small habitat patches, though individually species-poorer, often collectively enhance gamma diversity by supporting distinct species assemblages (i.e., higher beta diversity), especially when they are environmentally heterogeneous (Fahrig, 2020; Perrin et al., 2025; Riva & Fahrig, 2022). Small patches may further sustain metacommunities by functioning as dispersal hubs, stepping-stones, or buffers against environmental stochasticity (Altermatt & Ebert, 2010; Luo et al., 2022; Wang & Altermatt, 2019). Nevertheless, the value of small patches is not universal. When patches become excessively small, negative edge effects, resource scarcity, and demographic instability may limit their ability to support viable populations, especially when isolated from other patches (Deane & Riva, 2025; Fahrig, 2020; Lomolino, 2000).

Despite limited empirical evidence for minimum area constraints, habitat patches below area thresholds (e.g., <100 ha) are often perceived as having low conservation value (Riva & Fahrig, 2023b). In cities, however, the vast majority of habitat patches are much smaller (e.g., <1–2 ha; Kabisch and Haase 2014; Egerer et al. 2024). Although urban habitat patches are frequently of low quality (e.g., pathways, lawns; Nordh et al. 2009), characterized by high perimeter-area ratios (e.g., narrow, elongated shapes), and embedded within a difficult-to-navigate matrix, they nonetheless form heterogeneous mosaics that support diverse communities, often distinct from those in more natural ecosystems (Alberti et al., 2020; Aronson et al., 2016; Perrelet et al., 2024). Most urban biodiversity guidelines continue to emphasize minimum habitat areas ranging from 4.4 to >50 ha, based on taxa-specific sensitivity to urbanization (Beninde et al., 2015; Donnelly & Marzluff, 2004). However, these thresholds very commonly exceed the size of most urban green spaces (Gavrilidis et al., 2022) and are largely derived from patch-level inference (i.e., assuming species richness scales with area), overlooking the contribution to beta and gamma diversity of small patches. A macroecological perspective on cities, recently identified as a major barrier to effective urban biodiversity management (Casanelles-Abella et al., 2025), is therefore needed to develop guidelines that better account for landscape-level biodiversity contributions.

Despite uncertainties about how the area and spatial configuration of urban green spaces affect citywide biodiversity, growing evidence from the SLOSS literature shows that small urban green spaces, even <1 ha, can support substantial biodiversity, including plants (Godefroid & Koedam, 2003; Vega & Küffer, 2021), butterflies (Lizee et al., 2016), and birds (La Sorte et al., 2023). While these studies highlight the value of certain small patches (e.g., parks) for specific taxa, broader generalizations across habitat types, quality, and taxonomic groups are needed to reassess urban biodiversity guidelines. Vertebrates, for instance, may require larger patches due to greater biomass and energetic demands yet traverse urban matrices more easily than invertebrates (Moreno & Teixido, 2025; Prugh et al., 2008). Vertebrates and invertebrates may also differ in their responses to environmental factors (e.g., vegetation structural complexity), in both magnitude and spatial scale (Dietzel et al., 2024; Threlfall et al., 2017), thereby influencing how patch quality interacts with area to shape biodiversity patterns (Wang & Altermatt, 2019). In contrast, urban trees are largely decoupled from traditional ecological constraints, with distributions shaped by planting and management rather than dispersal or competition (Aronson et al., 2016; Avolio et al., 2021), and are thus expected to show weaker or different associations with patch area and connectivity than fauna. Without multi-taxon, spatially explicit monitoring that integrates habitat characteristics and environmental context across urban green spaces, the relative biodiversity contributions of small versus large patches remain poorly understood.

In this study, we examined the alpha, beta, and gamma diversity contributions of habitat patches of different areas for multiple taxonomic groups in an urban context. Leveraging a decade of observational data collected across the city of Zurich, Switzerland, we evaluated the contribution of habitat patch area and associated environmental variables to the biodiversity of invertebrates, vertebrates, and trees (Fig. 1). Specifically, we addressed three key questions: (1) How much do smaller patches contribute to biodiversity in comparison to larger patches? (2) Are these patterns consistent across taxonomic groups? (3) What roles do environmental factors and human management play in shaping biodiversity patterns across taxonomic groups and the patch area gradient? By addressing these questions, we aim to refine our understanding of fragmentation effects in urban ecosystems and inform the design of urban landscapes for biodiversity conservation.

**Figure 1:**
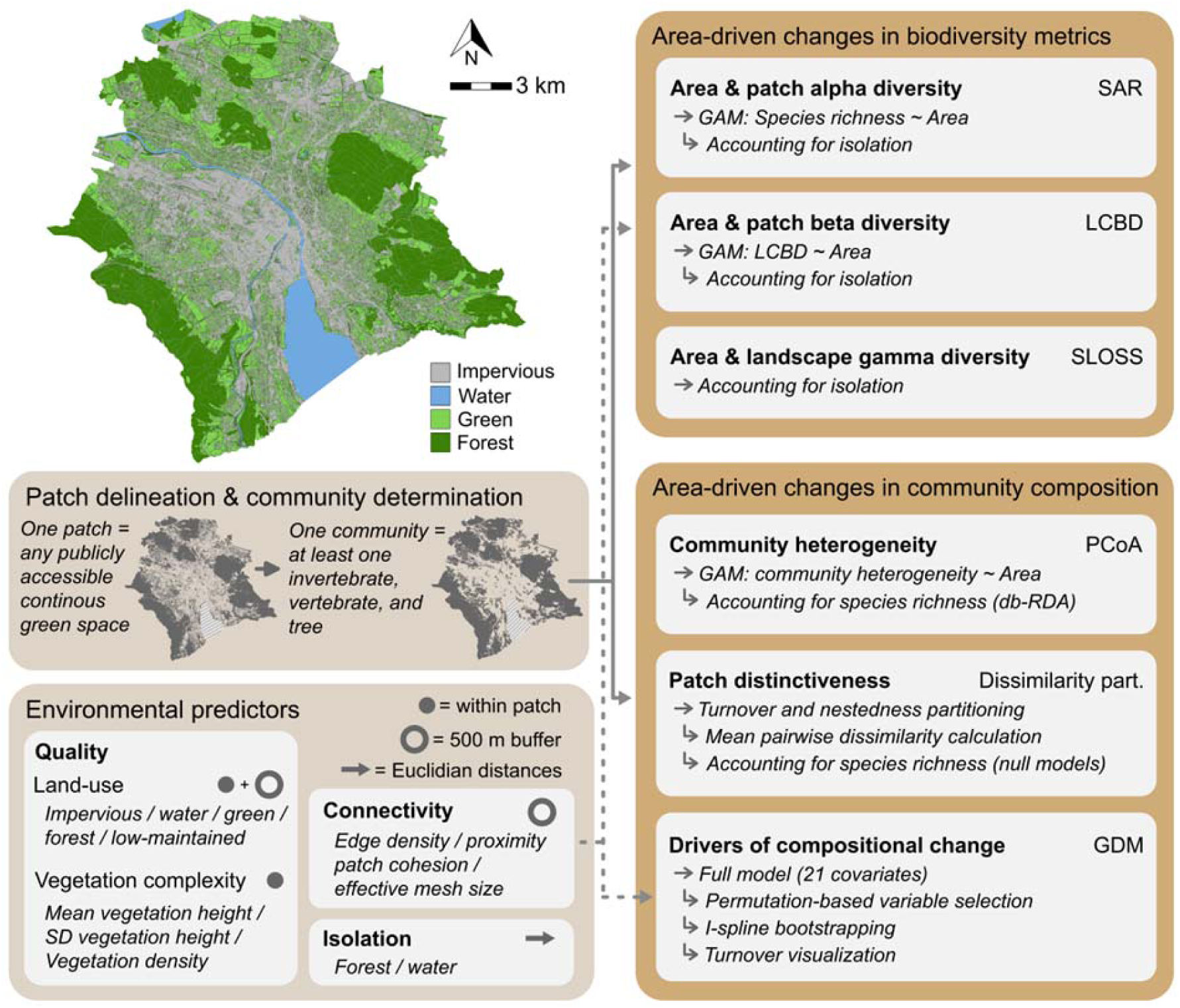
Study area (top right) and methodological workflow. Left: data processing (patch delineation, community identification, predictor extraction) with symbols indicating spatial scale: full circles = within-patch, empty circles = 500 m buffer from patch margins, arrows = shortest distances to forested habitats or water bodies. (Right: modelling framework assessing effects of patch area on biodiversity metrics and community composition, with individual model goals in bold and intermediate steps in italics. SAR = species-area relationship; LCBD = local contribution to beta diversity; SLOSS = single large or several small; PCoA = principal coordinate analysis; GDM = generalized dissimilarity modeling.

## Methods

### Study area and patch delineation

This study was conducted in Zurich, Switzerland’s largest city (total municipal area of 92 km^2^; 47°22′N, 8°33′E). Zurich is a mid-sized European city with ∼450,000 inhabitants (2024 census), undergoing relatively rapid urban expansion (United Nations, 2018). Densely built, the city includes extensive forested habitats (∼21 km^2^; 23% of total area) and a variety of other publicly accessible green spaces (∼29 km^2^; 31.5%), ranging from urban parks to unmanaged meadows (Fig. 1).

To quantify habitat fragmentation, we delineated habitat patches as all publicly accessible green spaces, excluding impervious surfaces and private gardens, based on a high-resolution green space map (Stadt Zürich, 2021). Although ecologically relevant (Goddard et al., 2010), private gardens were excluded due to limited biodiversity observations (see *Community determination*) and small share of total green space (∼10%). Contiguous green spaces were treated as single habitat patches, ignoring discontinuities ≤2 m (e.g., footpaths). This threshold was selected because larger gaps (e.g., 10 m, typical road width; Barthelmess, 2014) would imply mortality barriers, whereas smaller ones (e.g., 0.5 m) would artificially fragment continuous green spaces (Barthelmess, 2014). Sensitivity analyses indicated that results were robust to ±1 m variation in the threshold. Patch area was calculated for each delineated unit and log-transformed prior to statistical analyses, although values are also reported in hectares for interpretability.

### Community determination

Species occurrence data were compiled from a city-wide biodiversity monitoring program conducted between 2008 and 2017, during which a different urban district was surveyed each year, ensuring full spatial coverage of publicly accessible green spaces (excluding private gardens, rooftops, and railways). Although community composition can change over short time scales (Altermatt, 2012), we assumed that temporal turnover within the sampling period was small relative to the effects of habitat area and configuration (Uchida et al., 2018). The dataset included vertebrates from the Clade Tetrapoda (birds, mammals, reptiles, and amphibians), invertebrates from the Classes Insecta, Arachnida, and Gastropoda (butterflies, grasshoppers, dragonflies, beetles, true bugs, wasps, bees, spiders, and snails), and trees (Clades Angiosperms and Gymnosperms), all identified to species level. Trees were included for two reasons: they may mediate SLOSS patterns in other taxa (e.g., providing nesting resources for birds; Threlfall et al. 2017) and, because their distributions in cities largely reflect planting decisions rather than natural demographic processes (Aronson et al., 2017; Avolio et al., 2021), they offer a useful contrast to faunal taxa more directly governed by colonization-extinction dynamics. We did not distinguish between native and non-native species, as—for faunal groups—non-native species represented only a small fraction of recorded species, and—for trees—exploratory analyses indicated that accounting for tree origin had negligible effects on the results (Fig. S1). We therefore consider all taxa irrespective of origin for consistency.

Occurrences outside delineated habitat patches (<20% of records) were excluded, as they may result from unsystematic observations from private properties or temporary use of non-habitat areas (e.g., rooftops). Local ecological communities were defined as binary, presence-absence assemblages within each patch and analyzed separately for vertebrates, invertebrates, and trees (Fig. 1). While representing only a coarse classification, this taxonomic separation captures key ecological differences in dispersal ability, trophic ecology, and biomass, while ensuring adequate sample size for comparative analyses.

To ensure comparability across taxa, analyses were restricted to patches containing at least one species from each group, yielding 452 patches (Fig. S2). Very small patches (<0.1 ha) were most frequently excluded, primarily due to fewer vertebrate and invertebrate observations (Fig. S3). Although patch distribution was skewed toward the urban periphery, retained patches spanned a broad gradient of urban densification intensity, measured as the fraction of impervious surface cover within a 500 m buffer from patch margins (Fig. S4). Efforts were taken throughout all following analyses to account for uneven patch distribution (see *Area-based biodiversity contributions*). Given the 10-year duration, the high and well-distributed number of observations across the city, and the strong increase in species observations with patch area (Fig. S5), we assumed high sampling completeness across patches, minimizing area-related sampling artefacts (Fahrig, 2020).

### Environmental predictors

Although patch area was our primary focus, we also accounted for variation in habitat quality, connectivity, and isolation using a suite of 19 environmental covariates known to influence urban biodiversity patterns (Casanelles-Abella et al., 2021; Dietzel et al., 2024) (Fig. 1). Habitat quality was characterized by land-cover composition, including the fractions of impervious surfaces, water, and green spaces (further distinguishing forest and extensively maintained vegetation), quantified both within each patch and within a 500 m buffer surrounding patch margins. Habitat quality was further characterized by LiDAR-derived metrics within each patch, including mean and standard deviation of vegetation height (a proxy for structural complexity) and vegetation density. Habitat connectivity was described by four different metrics, including: edge density, mean proximity to neighboring patches (henceforth, “proximity), patch cohesion, and effective mesh size (for definitions, see *Supplementary Material I*), all calculated within a 500 m buffer of each patch’s margins, following Dietzel et al. (2024). Isolation from potential source habitats (Hanski, 1998) was assessed as the Euclidean distance from each patch edge to the nearest forest and permanent water body. Elevation was excluded due to its limited variation across the study area (maximum ∼450 m). Further details on environmental data extraction are provided in *Supplementary Material I*.

To visualize environmental heterogeneity among patches, we performed a Principal Component Analysis (PCA) on the measured environmental variables and overlaid smooth Generalized Additive Models (GAM) surfaces to examine how variation aligned with patch area.

### Area-based biodiversity contributions

To assess how patch area contributes to biodiversity patterns, we quantified its effects on alpha, beta, and gamma diversity for each taxonomic group (Fig. 1).

Patch-level (alpha) diversity was analyzed by fitting GAMs of log-transformed species richness against log-transformed patch area (Gaussian distribution, k = 4), thus using SAR analysis. Because patch area and spatial distribution covaried (small patches predominated in the city center), isolation from large, peri-urban forests was included to account for potential confounding effects.

Patch contributions to beta diversity were quantified using Local Contribution to Beta Diversity (LCBD; Legendre & De Cáceres, 2013), which measures the compositional uniqueness of each patch relative to the city-wide species pool. We tested the significance of LCBD values using permutation tests (9,999 permutations) and modeled the relationship between LCBD and patch area using GAM (Gaussian distribution, k = 4). We also used binomial generalized linear models (GLMs) to assess whether the probability of a patch making a statistically significant contribution to beta diversity varied with area. Isolation from forests was again included as a covariate in all models to address potential confounding between patch area and proximity to the city center.

Lastly, contributions at the landscape scale (gamma diversity) were determined through a SLOSS analysis following Quinn & Harrison (1988). Specifically, we computed cumulative species-area curves by sequentially aggregating patches in ascending (small-to-large) or descending (large-to-small) order of area. We also calculated the SLOSS index, defined as the ratio of the area under the small-to-large and large-to-small curves, where values above one suggest a greater conservation value of multiple small patches relative to a single large one. Unlike other analyses in this study, patch area was not log-transformed in the SLOSS analysis, reflecting conventional SLOSS implementations in the literature (e.g., Fahrig 2020; Riva and Fahrig 2023a). To account for the uneven distribution of patches, we repeated the SLOSS analysis within subsets of patches grouped by the first, second, and third quantiles of distance to the nearest forest, representing close, intermediate, and far distances, respectively.

### Area-driven changes in community composition

To evaluate the influence of patch area on community composition, we conducted analyses at three complementary levels of biological complexity, capturing the effects of area on (i) *overall community heterogeneity*, (ii) *patch distinctiveness*, and (iii) *compositional change relative to other environmental drivers*. All analyses were performed separately for vertebrate, invertebrate, and tree communities.

First, *overall community heterogeneity* was assessed using Principal Coordinates Analysis (PCoA) based on Jaccard dissimilarities, with GAM-derived smooth surfaces overlaid to visualize how compositional gradients aligned with patch area. Because Jaccard dissimilarities are influenced by richness differences, we used distance-based redundancy analysis (db-RDA) to assess the unique and shared contribution of patch area and species richness.

Second, to quantify *patch distinctiveness*, pairwise Jaccard dissimilarities were partitioned into turnover and nestedness components. For each patch, mean turnover and nestedness were calculated across all pairwise comparisons and modeled as functions of log-transformed patch area using GAMs. To control for richness effects, observed patterns were compared with null expectations generated from 500 randomized community matrices in which species identities were randomly permuted across patches while maintaining species richness. Further details on dissimilarity partitioning are provided in *Supplementary Material II*.

Third, to assess the contribution of patch area to *compositional change relative to other environmental drivers*, we used generalized dissimilarity models (GDMs; Ferrier et al. 2007), following Mokany et al. (2022). GDMs relate pairwise community dissimilarities to environmental and geographic distances using I-spline transformations, capturing nonlinear, monotonic relationships and estimating partial ecological distances—i.e., the proportion of community change attributable to each predictor (Ferrier et al., 2007). For each individual group, only patches containing five or more species were included to avoid extreme dissimilarity values (e.g., 0, 0.5, 1) associated with depauperate assemblages.

For each group, a full model incorporating the previously mentioned 19 environmental predictors, log-transformed patch area, and geographic distances between patches (21 covariates total) was initially fitted. Model significance was evaluated by permuting all predictor values to confirm that the full model explained more variation than expected by chance. Predictor importance was quantified as the percent reduction in deviance explained when individual predictors were permuted, with significance determined from bootstrapped *p*-values. Non-significant predictors were iteratively removed via backward elimination, reassessing model significance at each step.

We then built a refined model using only predictors considered to be significant or marginally significant (p<0.1). Predictor correlations were generally low-to-moderate (maximum Pearson’s r = 0.53; Fig. S6). Model performance was evaluated from the correlation between observed and predicted ecological distances (Fig. S7). To evaluate robustness, refined models were refitted to 500 bootstrap subsets (90% of patches each), and distributions of I-spline functions were summarized to characterize the consistency of each predictor’s contribution to partial ecological distances. Finally, to determine spatial gradients in predicted community turnover, we projected GDM-transformed ecological distances for all surveyed patches. For visualization purposes, as compositional change is multidimensional, we used a PCA to reduce its dimensionality, using the first principal component to represent the dominant axis of variation.

All analyses were conducted using R v4.5.0 (R Core Team, 2025). Key packages used are reported in Table S1.

## Results

We surveyed 452 urban habitat patches (median area = 1.4 ha; min-max = <0.05–761 ha; Fig. 4) to assess biodiversity patterns across three major taxonomic groups: invertebrates (488 species), vertebrates (166 species), and trees (416 species) (Figs. 1 and 2). For all groups, species richness distributions were highly skewed, with a few species-rich (typically large, i.e., ≥5 ha) patches and many species-poor (typically small, i.e., <5 ha) patches (Fig. S8).

### Area-based biodiversity contributions

All taxa exhibited a positive SAR, although this value showed signs of plateauing at both ends of the area gradient for faunal groups (i.e., triphasic shape), especially invertebrates (Fig. 2A). Small patches also contributed disproportionately to beta diversity, with LCBD generally declining as patch area increased before rising again in very large patches—except for trees, where this later increase was weaker—and almost never returning to the highest, significant LCBD values observed for all taxa (Fig. 2B). Importantly, isolation from forests had non-significant to weak effects on both alpha and beta diversity patterns relative to patch area (Tables S2 and S3). For all taxa, small habitat patches were disproportionately more likely to contribute significantly to beta diversity (GLM; p <0.001; Fig. S9; see triangles in Fig. 2, second row). In fact, patches in the smallest area quartile had a 43%–47% chance of significantly contributing to beta diversity, compared to <0%–7% for those in the largest quartile, depending on taxon (Fig. S10). SLOSS analyses further indicated that, per unit area, small patches cumulatively supported more species than equivalent aggregations of large patches, with the strongest effect for trees, followed by invertebrates and vertebrates, for which the benefit of small patches was less conclusive (i.e., two lines crossing; Fig. 2C). Stratifying SLOSS analyses by isolation from forests produced qualitatively similar results, indicating that the gamma diversity contribution of small patches was largely independent of isolation (Fig. S11).

**Figure 2:**
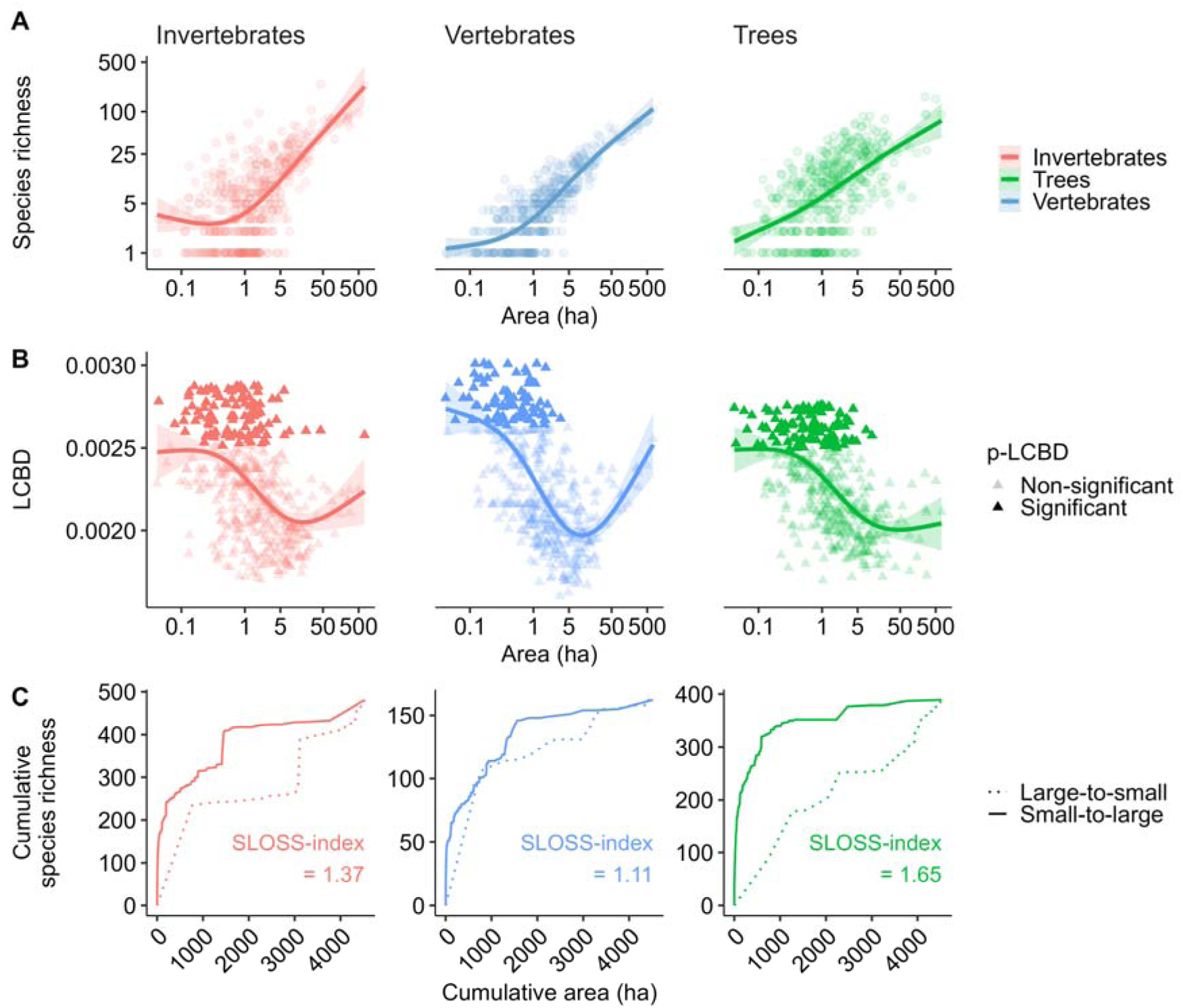
Contribution of small habitat patches to invertebrate, vertebrate, and tree biodiversity (color-coded) to (A) alpha, (B) beta, and (C) gamma diversity. (A) GAM (k = 3) regressions between log-transformed patch area and log-transformed species richness, with each dot representing a site and color-coded by taxonomic group. (B) GAM (k = 3) showing relationship between patch area (log-scaled) and Local Contribution to Beta Diversity (LCBD); shade indicates LCBD significance, ribbon shows standard error of model predictions. (C) SLOSS analysis showing cumulative species richness when adding small (solid) vs. large (dashed) patches first; SLOSS-index >1 indicates stronger contribution from small patches.

### Area-driven changes in community composition

Both GDM analyses and PCoA ordinations highlight strong area effects on community composition (Fig. 3 and S12). Small patches supported highly divergent assemblages dominated by turnover, whereas large patches harbored more homogeneous assemblages characterized by nestedness (Fig. 4 and S12). These differences were not simply attributable to richness: db-RDA, which relates community dissimilarities to explanatory variables, showed that patch area explained a similar portion of the compositional variation as richness alone (R^2^ ≈ 0.05–0.08; Fig. S13). Similarly, observed patterns of turnover and nestedness deviated from null expectations (Fig. S12 and S14).

**Figure 3:**
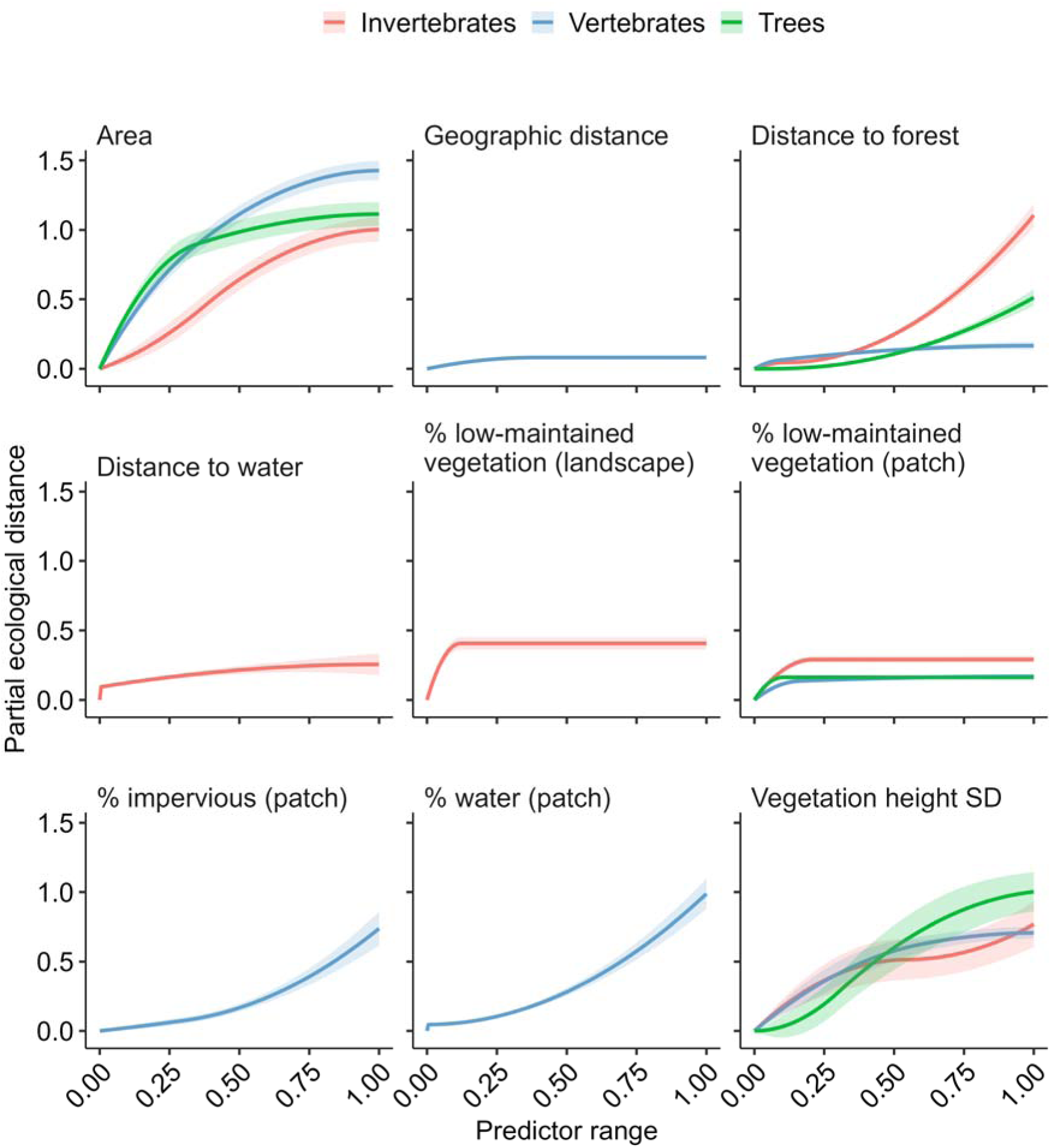
I-spline functions (±90% CI) illustrating the effects of significant predictors (scaled 0–1) on partial ecological distance for invertebrate, vertebrate, and tree communities (color-coded). Non-significant predictors are omitted; absence of a line indicates a predictor was non-significant for that taxonomic group. Curves show median fits across 500 bootstrap iterations using random subsets comprising 90% of the sites.

**Figure 4:**
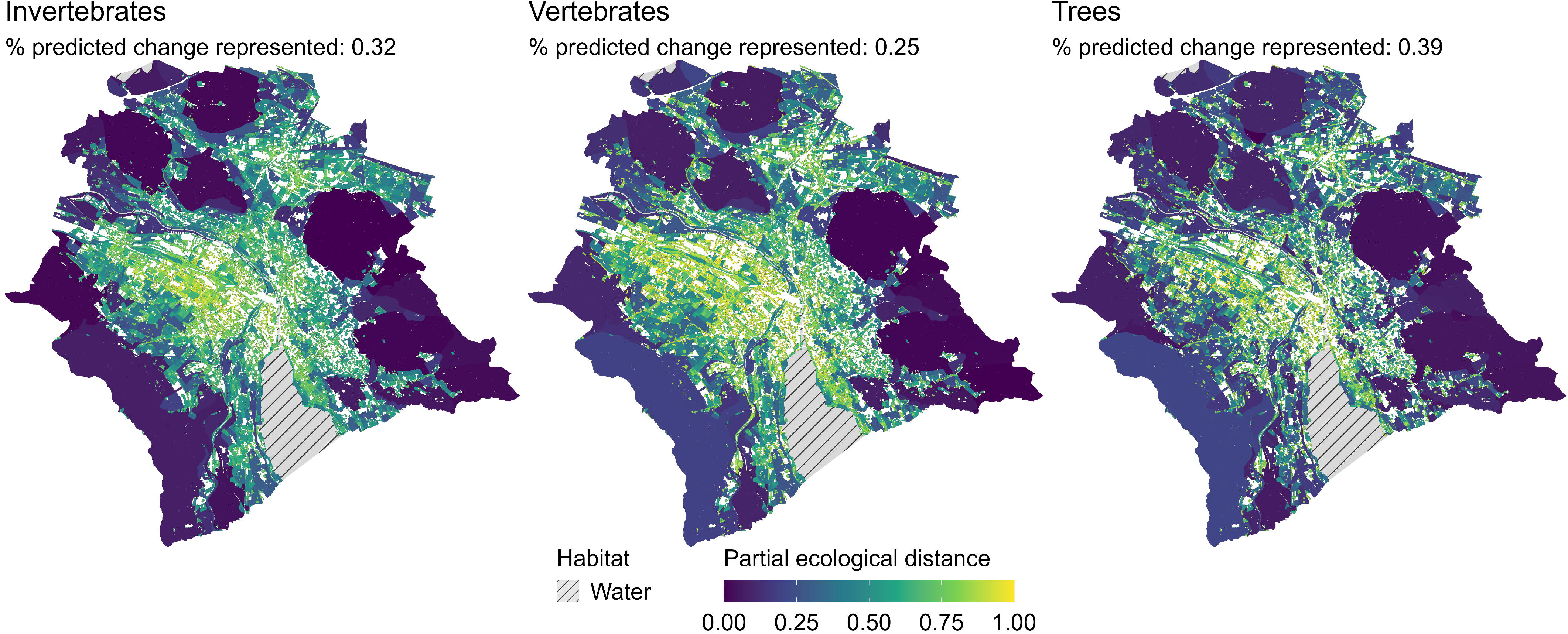
Spatial patterns of predicted community change, represented by the first principal component (PC1) from a PCA on GDM-predicted partial ecological distances (scaled 0–1). Percentages of explained variance of predicted change are indicated in the subtitles. Hatched areas denote excluded water patches.

Small habitat patches exhibited greater heterogeneity than larger patches in their environmental conditions (Fig. S15). In fact, GDM confirmed that these environmental factors played an important role in determining community composition, jointly with patch area and geographic location, with the strength of these effects varying across taxa (Fig. 3). Deviance explained was moderate for invertebrates and vertebrates (33.7 and 34.9%, respectively) and lower for trees (15.2%) (Fig. S16). Patch area emerged as a consistent predictor across groups, explaining 11.9% of deviance for vertebrates, but only 6.8% and 4.9% for invertebrates and trees, respectively, with much of its influence mediated through interactions with environmental variables (up to 11.5% and 5.2%, respectively; Fig. S16). Importantly, these results show that patch area was associated with shifts in community composition within patches, which is not inconsistent with the complementarity of patches in a landscape tested by the SLOSS analysis.

Among 19 environmental predictors (Fig. 1), the most frequently selected were isolation from forests, related to vegetation complexity (low-maintenance cover, local variation in vegetation height), and water availability (isolation from water, water cover; Fig. 3). Isolation from forests exerted particularly strong effects on invertebrates, while vegetation complexity consistently influenced all groups. Both faunal groups also responded to either presence (i.e., patch-level cover) or isolation from water bodies. Idiosyncratic responses include invertebrate communities being sensitive to landscape-scale vegetation maintenance, and vertebrate communities responding to local impervious surface cover. Geographic location was only important for vertebrates. Because patch area was negatively correlated with distance to forest (Fig. S4), area and environmental factors jointly led to coherent patterns of compositional change across taxa (Fig. 4). Core urban areas, characterized by small, isolated patches, emerged as hotspots of predicted community change, especially for trees, while peripheral, larger patches closer to forest showed greater similarity. Although broad patterns were consistent across taxa, finer-scale differences reflected taxon-specific sensitivities: invertebrate communities exhibited a slightly more gradual shift, whereas vertebrate and tree assemblages showed patchier, mosaic-like patterns of change (Fig. 4).

## Discussion

Our study provides a comprehensive, multi-taxon perspective on how patch area shapes urban biodiversity. As expected from the species-area relationship (MacArthur & Wilson, 1967), larger patches generally supported higher local (alpha) richness. However, smaller patches (i.e., 0.1–5 ha) contributed disproportionately to compositional (beta) and regional (gamma) diversity, with patterns broadly consistent across taxa. Patch area was also consistently associated with differences in community composition, with the strength and direction of these relationships mediated by taxon-specific environmental sensitivities.

Despite lower species richness, small patches supported distinct assemblages that contributed disproportionately to compositional turnover, even after accounting for uneven spatial distribution (e.g., isolation from forests). In fact, they were approximately six times more likely than large patches to significantly contribute to beta diversity. SLOSS analyses further indicated that, taken together, groups of small patches supported more species per unit area than single large patches. While several mechanisms could underlie these patterns, including patch age, clustered species distributions, or elevated immigration rates (Fahrig, 2020), our results suggest high environmental heterogeneity is the dominant driver, potentially reflecting the widespread distribution of small patches compared with the peri-urban concentration of large patches. In fact, environmental heterogeneity may be weaker in large urban patches, as they are often intensively managed for specific human uses (e.g., picnics) by centralized administrators or service companies, whereas small patches may be more neglected or managed by diverse actors with heterogeneous practices (Aronson et al., 2017; Donati et al., 2025). Although the uneven distribution of patches in the city in relation to their area could also contribute to the observed patterns, the weak effect of isolation from forests relative to area on alpha and beta diversity, along with consistent SLOSS outcomes after stratifying by isolation, confirms that patch area is a strong determinant of beta and gamma diversity contributions. Collectively, these findings challenge conventional urban planning thresholds (e.g., >50 ha for urban avoider species, >4 ha for urban-tolerant species; Beninde et al., 2015), demonstrating that small patches, even below 1 ha, make essential contributions to urban biodiversity at the landscape scale (Riva & Fahrig, 2022, 2023b). We stress that this does not imply that habitat loss is inconsequential, but rather that remaining green spaces often retain high ecological value even when small, highlighting the importance of both their preservation and creation (e.g., through blue-green infrastructure) when possible. Similarly, while isolation effects were weak—potentially due to the limited geographic extent or high density of habitat patches in Zurich—connectivity should not be overlooked, and preserving and implementing small patches acting as sources or stepping stones can facilitate species persistence in fragmented landscapes (Altermatt & Ebert, 2010; Luo et al., 2022).

Beyond species richness, patch area strongly impacted community composition. Small patches supported distinct assemblages with higher species turnover, whereas large patches contained a greater but more homogeneous subset of the regional species pool, resulting in higher nestedness. These patterns could not be explained by richness alone, but emerged from the combined effects of area and environmental characteristics. When considered individually, patch area generally had a stronger and more linear influence on species composition than most environmental factors. Nonetheless, environmental characteristics, including vegetation complexity, water availability, and isolation from forests, exerted substantial influence, particularly at higher dissimilarity values and in combination with patch area. Because these drivers influenced taxa consistently, groups exhibited broadly parallel compositional shifts along the peri-urban to urban gradient. Large peri-urban patches, typically located on city outskirts with lower impervious cover and closer proximity to forests, hosted compositionally similar assemblages. In contrast, small patches, especially within urban cores but also near forested areas, exhibited heterogeneous management and environmental conditions, leading to communities that were both unique and spatially variable. This aligns with recent evidence that compositional turnover, rather than homogenization, predominates in anthropogenically modified landscapes (Keck et al., 2025).

Some taxon-specific responses also emerged. Tree communities were the most challenging to interpret, with strong SLOSS relationships driven by contrasting processes in large, relatively species-poor semi-natural forests versus smaller, horticulturally enriched patches shaped by human management (Aronson et al., 2017; Augustinus et al., 2024; Kendal et al., 2012). This imprint of horticulture was also evident in GDM results, where tree composition was best explained by patch area—indicative of planting opportunities—and vegetation complexity, a variable derived from trees themselves, introducing circularity in explanatory power (Aronson et al., 2017). Nonetheless, these results suggest that horticultural choices can enhance the contributions of small patches (Egerer et al., 2024), which may in turn benefit other taxa (e.g., birds; Threlfall et al., 2017).

For faunal groups, collections of small patches had a strong positive effect on invertebrates, though the effect was less pronounced for vertebrates. These findings likely reflect differences in mobility, metabolic and home-range requirements, and sensitivity to urban stressors such as noise and light pollution (Kunc & Schmidt, 2019; Moreno & Teixido, 2025; Prugh et al., 2008). For both groups, the interplay between patch area, quality, and isolation determined community composition. Invertebrate composition shifted primarily with landscape-scale factors, including isolation from forests, proximity to water bodies, and vegetation maintenance practices, whereas vertebrates responded more to local factors such as impervious cover and vegetation management. These differences may reflect dispersal ability, with low-mobility invertebrates showing gradual turnover and more mobile vertebrates exhibiting mosaic-like distributions (Braschler et al., 2020; Holtmann et al., 2017; Knop, 2016). As a result, it is important to maintain a diversity of patch sizes and habitat qualities, such as both high and low vegetation complexity, to accommodate the contrasting life-history traits and dispersal strategies found across taxa (Wang & Altermatt, 2019).

Despite the strengths of our multi-taxon, spatially explicit, city-wide dataset, some limitations of our analysis must be acknowledged. First, the use of presence-only data precluded assessments of demographic stability or genetic viability, limiting our ability to evaluate population persistence and genetic diversity, respectively, along the patch-area gradient (Fahrig et al., 2022; Fletcher et al., 2018; Multigner et al., 2025). Given the contribution of small patches to urban biodiversity, future work should determine whether such patches can sustain viable populations over extended periods, particularly when connectivity to larger, compositionally stable habitats is limited. If not, restoration to enlarge small patches may be required. Second, because larger patches tended to be mostly distributed at the city’s periphery, area and isolation from forests were not fully independent. Although isolation was included in all models, residual correlation may remain. Importantly, because the concentration of large patches in peri-urban areas is a common pattern in cities (e.g., Kabisch and Haase, 2014), our results reflect general compositional trends along urbanization gradients. Lastly, broad taxonomic groupings limited examination of finer functional or life-history traits (e.g., volant vs. non-volant, terrestrial vs. semi-aquatic taxa). While finer-scale groupings can provide ecological insights (e.g., Huber et al., 2025), they would have yielded low-diversity assemblages with unstable dissimilarity estimates (e.g., only 10 amphibian species in Zurich; Ineichen et al., 2022). We therefore focused on broader taxonomic groupings to maximize spatial coverage, retain high within-group richness, and maintain meaningful contrasts in dispersal, foraging strategies, and biomass. Future studies should explore finer trait- or clade-based approaches to better understand how patch area and landscape context influence phylogenetic and functional diversity in urban ecosystems (La Sorte et al., 2023).

## Conclusion

Our results demonstrate that while larger patches consistently supported higher local species richness, small patches made disproportionate contributions to beta and gamma diversity by harboring distinct assemblages that enhanced landscape-scale, cumulative biodiversity. These contributions were broadly consistent across taxa, although their magnitude and underlying drivers varied, potentially reflecting differences in dispersal mode and ability, habitat dependence, and sensitivity to urban stressors. Beyond the well-known relationship between species richness and area, we found that patch area was closely linked to community composition: small patches in urban cores supported unique communities shaped by high environmental heterogeneity within and around each patch, whereas large, peripheral patches harbored more nested and compositionally similar assemblages. Our results suggest that urban biodiversity in Zurich has not been homogenized but instead restructured along gradients of patch area, vegetation complexity, and landscape context (e.g., proximity to water and forested habitats). Conserving urban biodiversity, therefore, requires a dual strategy: protecting large, structurally complex patches while recognizing, preserving, and enhancing (e.g., through blue-green infrastructure implementation) the contributions of small patches. Because small patches are often the most threatened by urban development, their recognition as major contributors to regional biodiversity is essential for the conservation of urban-dwelling species.

## Acknowledgments

We thank the ETH Board for funding through the Blue-Green Biodiversity (BGB) Initiative (BGB 2021-2024).

## Data Availability Statement

Data and code to fit all models listed in this study are openly available on Zenodo with the identifier https://doi.org/10.5281/zenodo.18369652.

