## Supplementary Material for "Small habitat patches can be the largest contributors to urban biodiversity across taxonomic groups"

Kilian Perrelet,

Swiss Federal Research Institute for Forests, Snow, and Landscapes (WSL), Biodiversity and Conservation Biology, Zürcherstrasse 111, 8903 Birmensdorf, Switzerland

**Acknowledgments**

We thank the ETH Board for funding through the Blue-Green Biodiversity (BGB) Initiative (BGB 2021-2024).

**Conflict of Interest Statement**

The authors declare that they have no known competing financial interests or personal relationships that could have appeared to influence the work reported in this paper.

**Data Availability Statement**

Data and code to fit all models listed in this study are openly available on Zenodo with the identifier <https://doi.org/10.5281/zenodo.18369652>.

**Supplementary material**

I. *Environmental data extraction*

Although patch area was our primary variable of interest, we also accounted for a suite of environmental predictors known to shape urban biodiversity patterns (Beninde et al., 2015; Nielsen et al., 2014). In total, for each patch, we evaluated 19 variables capturing land-use and land cover (LULC) across local (within-patch) and landscape (500 m buffer from patch margin) scales, vegetation complexity, connectivity, and isolation.

- *Land-use and land cover*: We quantified LULC using a 10 m resolution raster derived from the official habitat map provided by the city of Zurich (Stadt Zürich, 2021). For each patch and its surrounding 500 m buffer—two spatial scales commonly used in urban biodiversity studies (Dietzel et al., 2024; Egerer et al., 2017; Nielsen et al., 2014; Philpott et al., 2014)—we extracted the proportions of impervious surfaces, blue spaces (water bodies), and green spaces. Green space was further subdivided into forest and low-maintenance green spaces (e.g., unmanaged meadows) based on a binary classification of habitat management intensity following Casanelles-Abella et al. (2021). All extractions were conducted using the *exactextractr* v0.10.0 (Baston & ISciences, 2023).
- *Vegetation structure*: We characterized vegetation complexity within patches using LiDAR-derived structural metrics, including vegetation density, mean vegetation height, and height variability, the latter serving as a proxy for vertical structural heterogeneity (e.g., Noordijk et al., 2010). These metrics were derived using the *lidaRtRee* package v4.0.8 (Monnet, 2023).
- *Connectivity*: We quantified connectivity using four widely applied metrics: edge density, proximity, patch cohesion, and effective mesh size, all calculated from the LULC raster with the *landscapemetrics* package v2.2.1 (Hesselbarth et al., 2019). The proximity index measures the mean distance to the nearest neighboring patch within a 500 m buffer. The patch cohesion index quantifies the degree of patch aggregation, ranging from 0 (all patches isolated) to 1 (all patches fully aggregated). Effective mesh size represents the average area of the patch in which a randomly selected point would fall, providing an interpretable measure of overall landscape connectivity or fragmentation (Jaeger, 2000). These metrics were selected because they are among the most commonly used indicators of connectivity in habitat-fragmentation research (Hanson et al., 2018). All metrics were calculated in a 500 m buffer from patch margins.
- *Isolation*: Finally, we quantified patch isolation as the Euclidean distance from each patch centroid to the nearest water body and nearest forest fragment, based on the LULC raster.

II. *Dissimilarity partitioning*

To assess whether patch area would drive changes in pairwise compositional dissimilarities, we partitioned Jaccard dissimilarity into species turnover and nestedness components. We then quantified for each habitat patch average pairwise Jaccard dissimilarity, turnover, and nestedness values by averaging across all pairwise comparisons involving the focal patch. We then used the same GAM analysis as before to assess the relationships between patch area and Jaccard dissimilarity, turnover, and nestedness, respectively. Finally, to assess whether observed patterns deviated from random assembly processes, we generated 500 null community matrices by randomizing species identities within patches while preserving observed richness and calculated for each the mean Jaccard dissimilarity, turnover, and nestedness to other patches across all randomized iterations. We also obtained the relative rate of changes by dividing the observed rate of change along patch area by the rate of change along patch area derived from null models. We then repeated these steps independently for each taxonomic group.

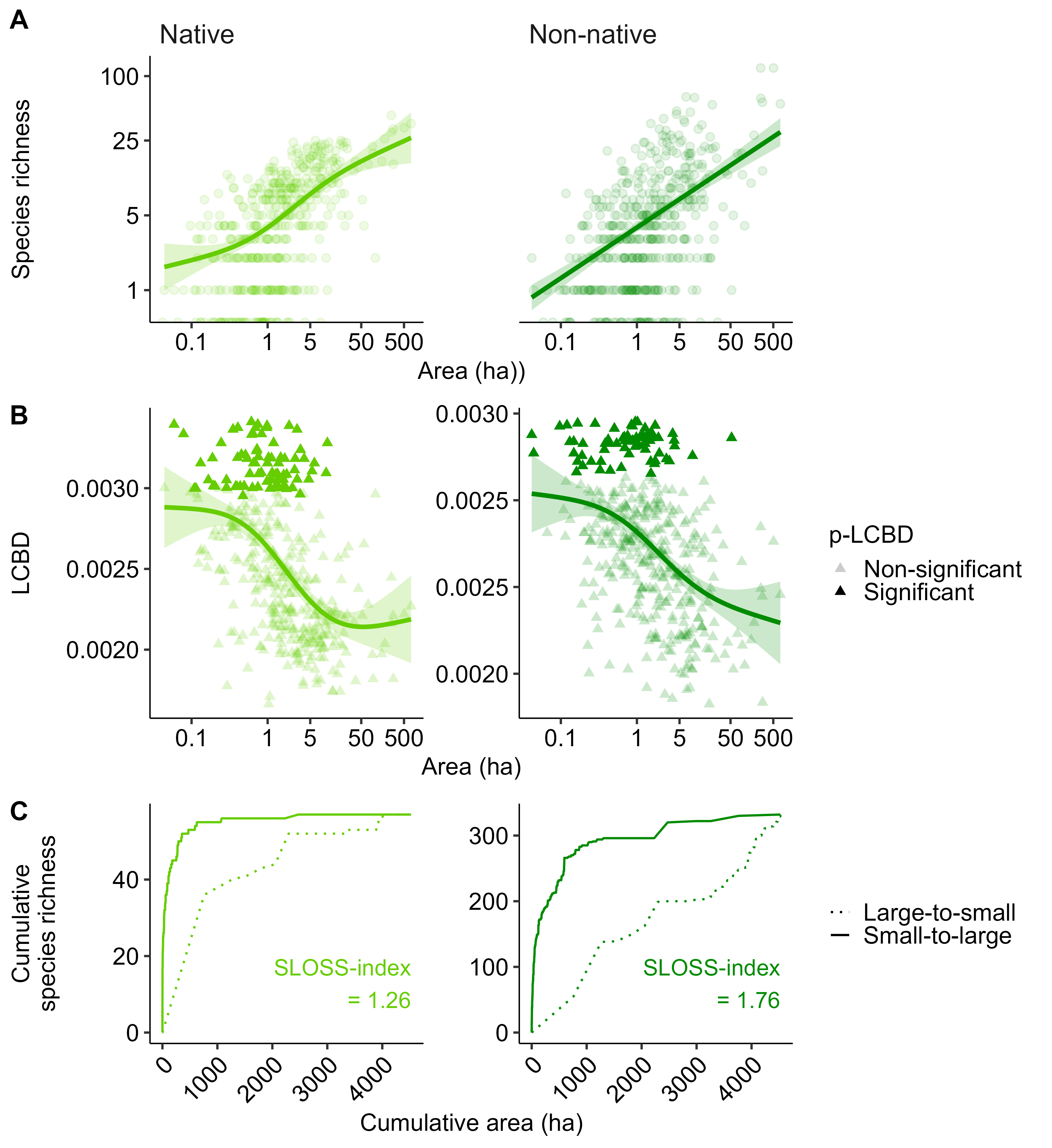
*Figure S1: Contribution of small habitat patches to tree biodiversity based on tree origin (native/non-native; color-coded) to (A) alpha, (B) beta, and (C) gamma diversity. (A) GAM (k = 3) regressions between log-transformed patch area and log-transformed species richness, with each dot representing a site and color-coded by taxonomic group. (B) GAM (k = 3) showing relationship between patch area (log-scaled) and Local Contribution to Beta Diversity (LCBD); shade indicates LCBD significance, ribbon shows standard error of model predictions. (C) SLOSS analysis showing cumulative species richness when adding small (solid) vs. large (dashed) patches first; SLOSS-index >1 indicates stronger contribution from small patches. patches.*

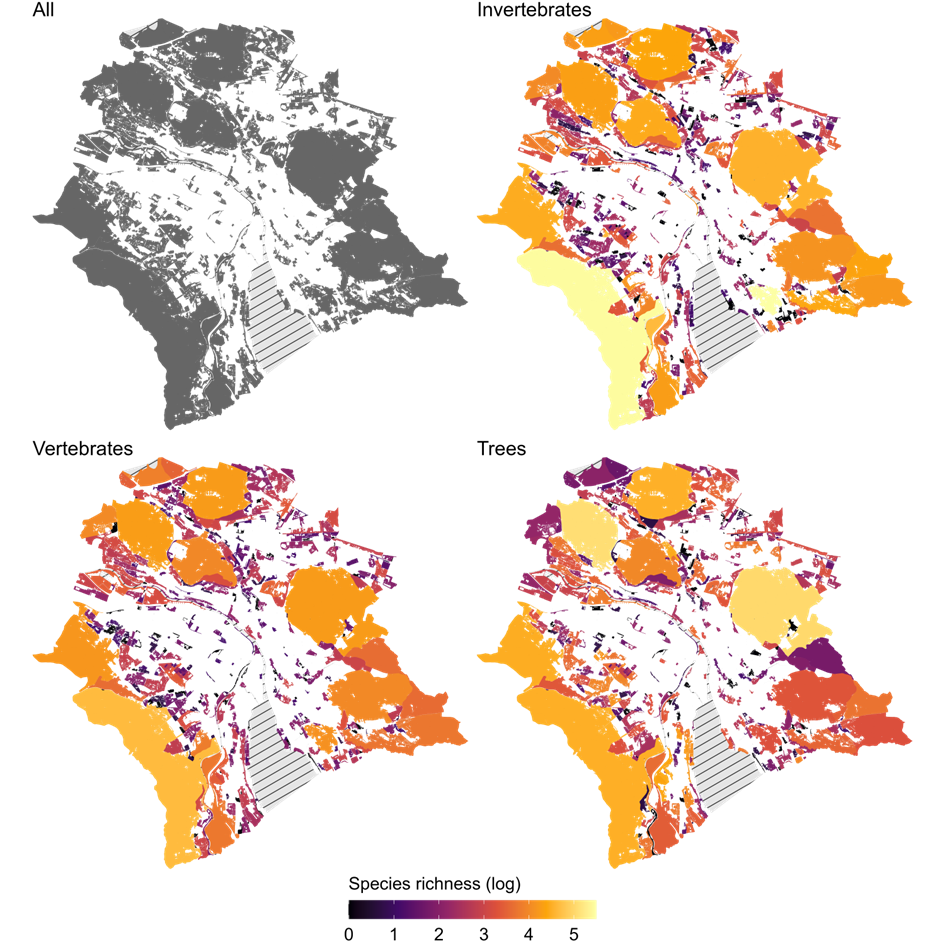
*Figure S2: Distribution of the 452 habitat patches considered. Maps show all considered patches, individually color-coded based on invertebrate, vertebrate, and tree species richness (log-transformed for visualization purposes). Hashed areas denote excluded water patches.*

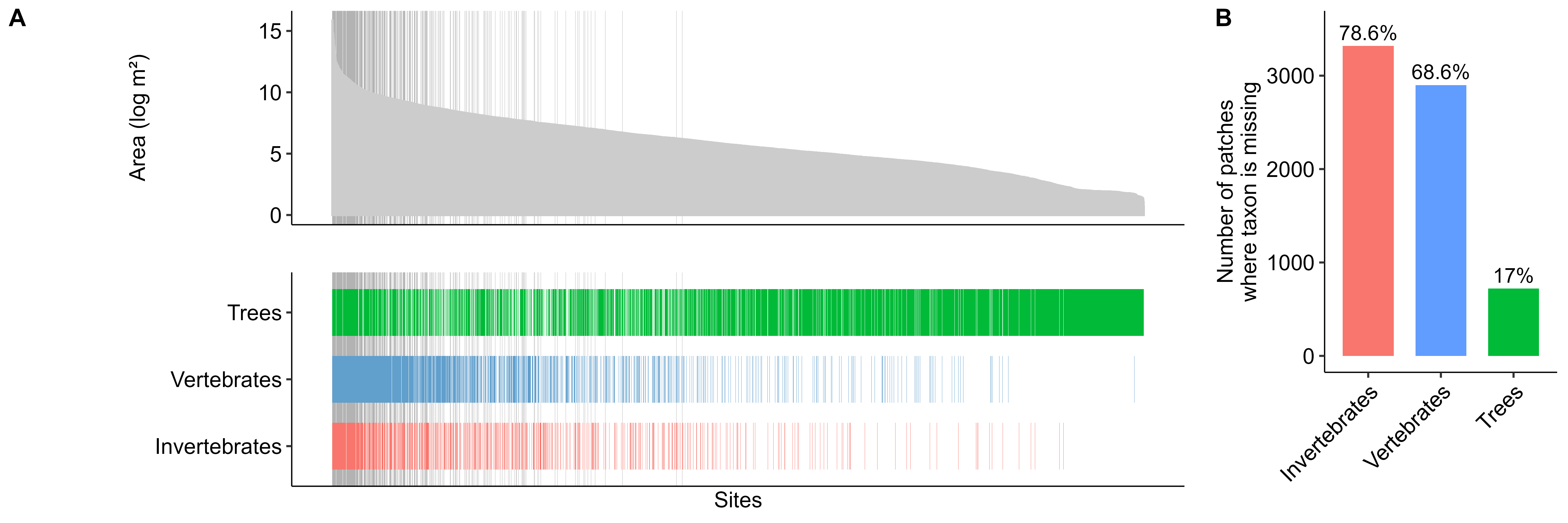
*Figure S3: Distribution of taxonomic groups across all potential habitat patches containing at least one species. (A) Patch area distribution (top) and patch-level presence of each taxonomic group (bottom). Bars indicate presence (color-coded by taxonomic group), with absent bars indicating the taxon was not detected. Sites are ordered by patch area. Thin grey lines highlight sites containing at least one species from each taxonomic group, which were retained for analysis. (B) Number of patches in which each taxonomic group was absent across all potential patches, color-coded by taxonomic group.*

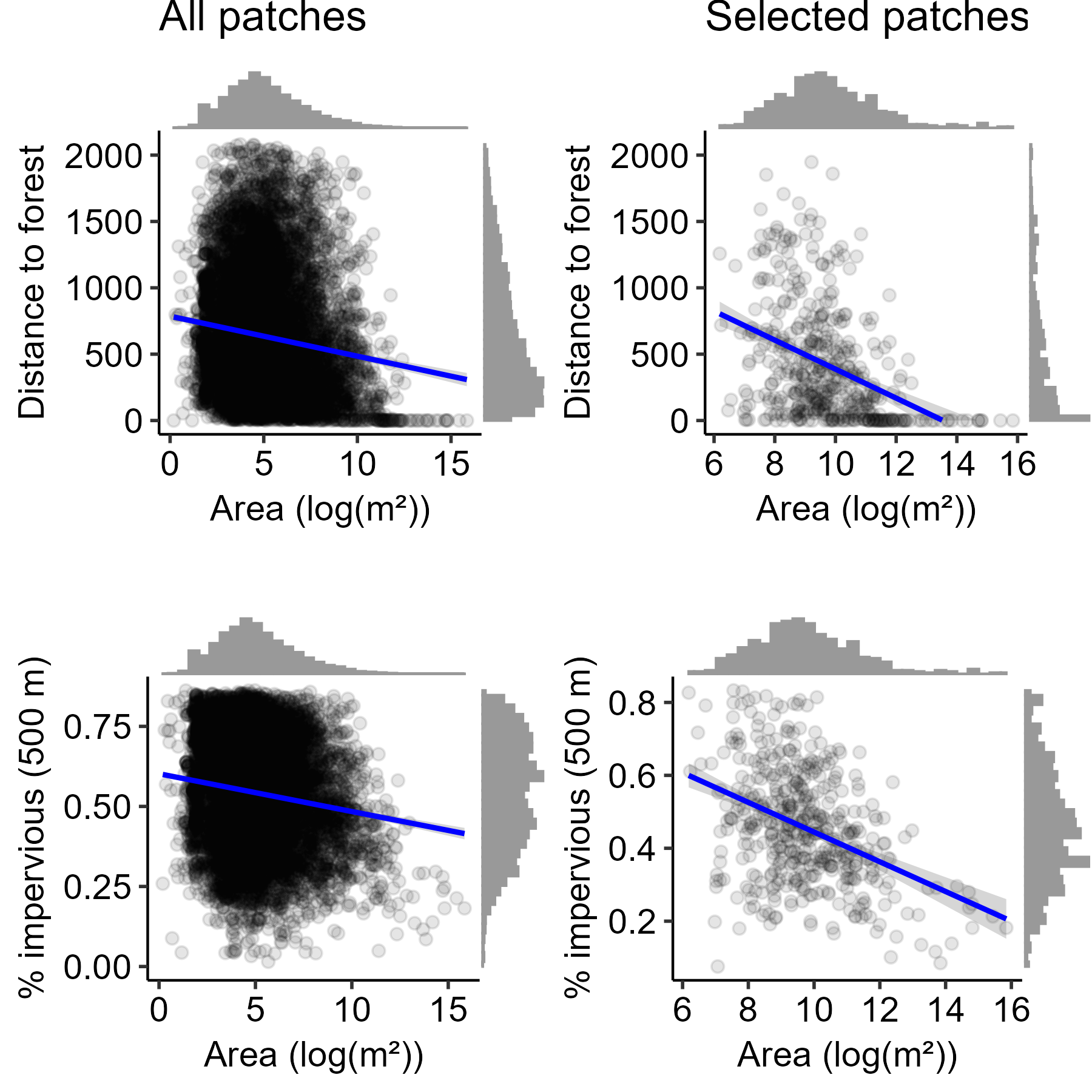
*Figure S4: Regression lines and histograms showing relationships between patch area (log-transformed) and distance to forest (top) and the proportion of impervious surface at a 500 m buffer (bottom), for all patches (regardless of the presence or absence of species) and those with ≥1 species of invertebrates, vertebrates, or trees (i.e., focal patches included in further analyses). Each dot represents a patch; all correlations are significant.*

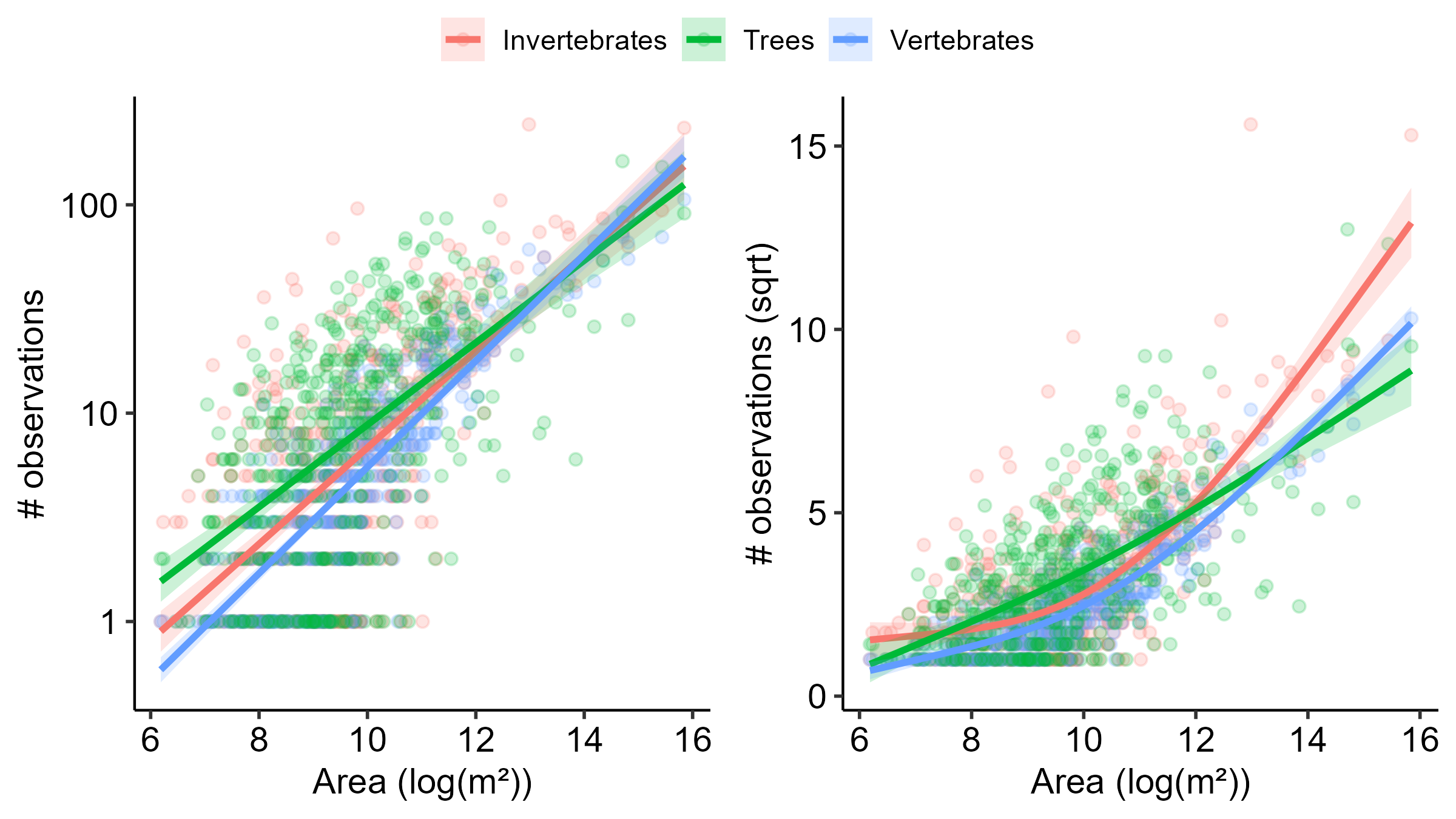
*Figure S5: GAM regressions between patch area (log-transformed) and total observations (right), and observations per m^2^ (log-transformed), for invertebrates, vertebrates, and trees (color-coded).*

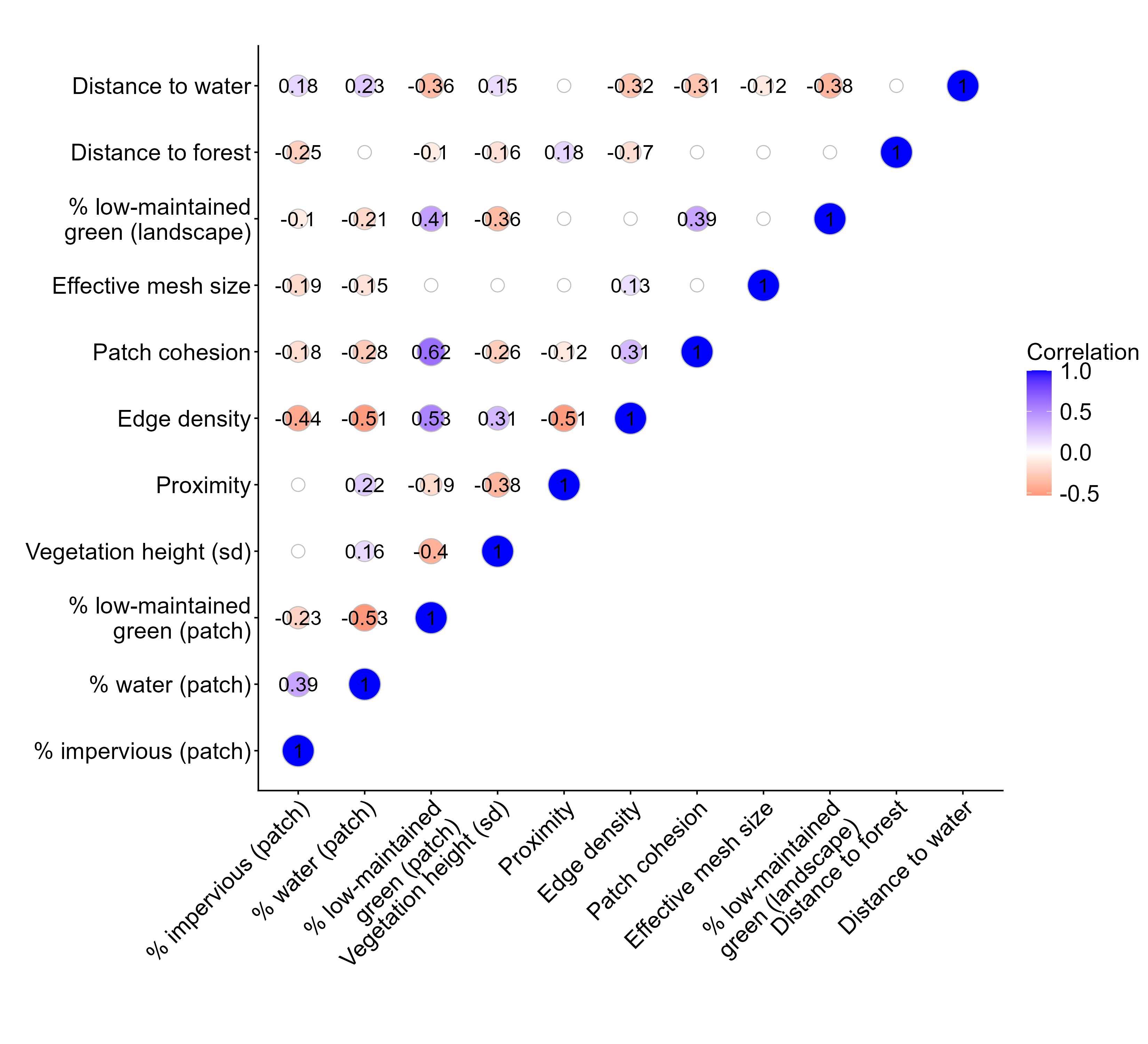
*Figure S6: Correlation matrix showing the correlation between covariates selected by the GDM framework, along with the connectivity metrics (edge density, proximity, patch cohesion, effective mesh size). White, unlabeled dots indicate non-significant correlations; labels and color represent significant correlations and their strength.*

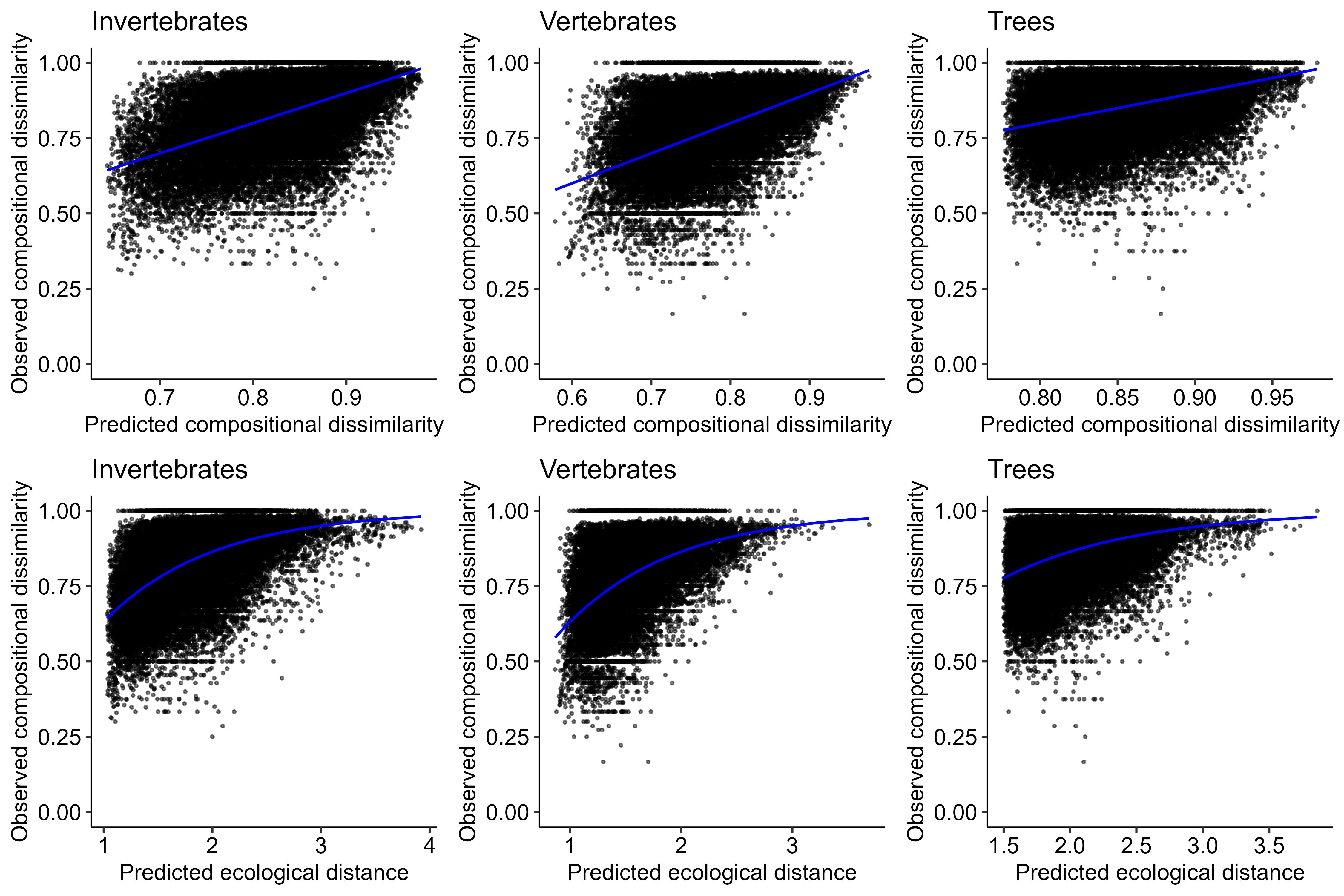
*Figure S7: Model diagnostic plots showing relationships between predicted versus observed compositional dissimilarity (top) and predicted ecological distance versus observed compositional dissimilarity (bottom) for invertebrate, vertebrate, and tree communities. Blue lines represent the fitted relationships from the GDM.*

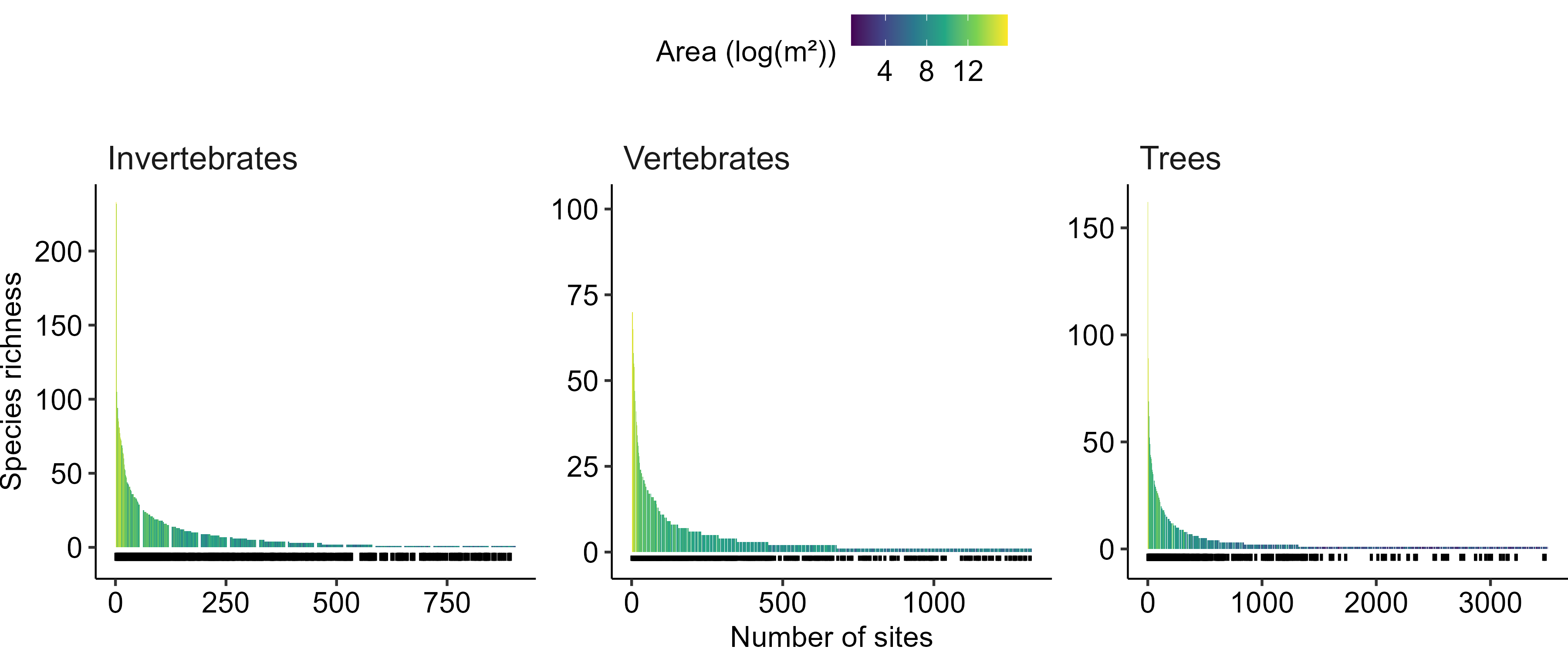
*Figure S8: Bar plot showing species richness across habitat patches for invertebrates, vertebrates, and trees. Each bar represents a patch, ranked from highest to lowest species richness within each taxonomic group, color-coded based on patch area (log-transformed). Black segments indicate sites retained in the analysis, containing at least one species from each taxonomic group.*

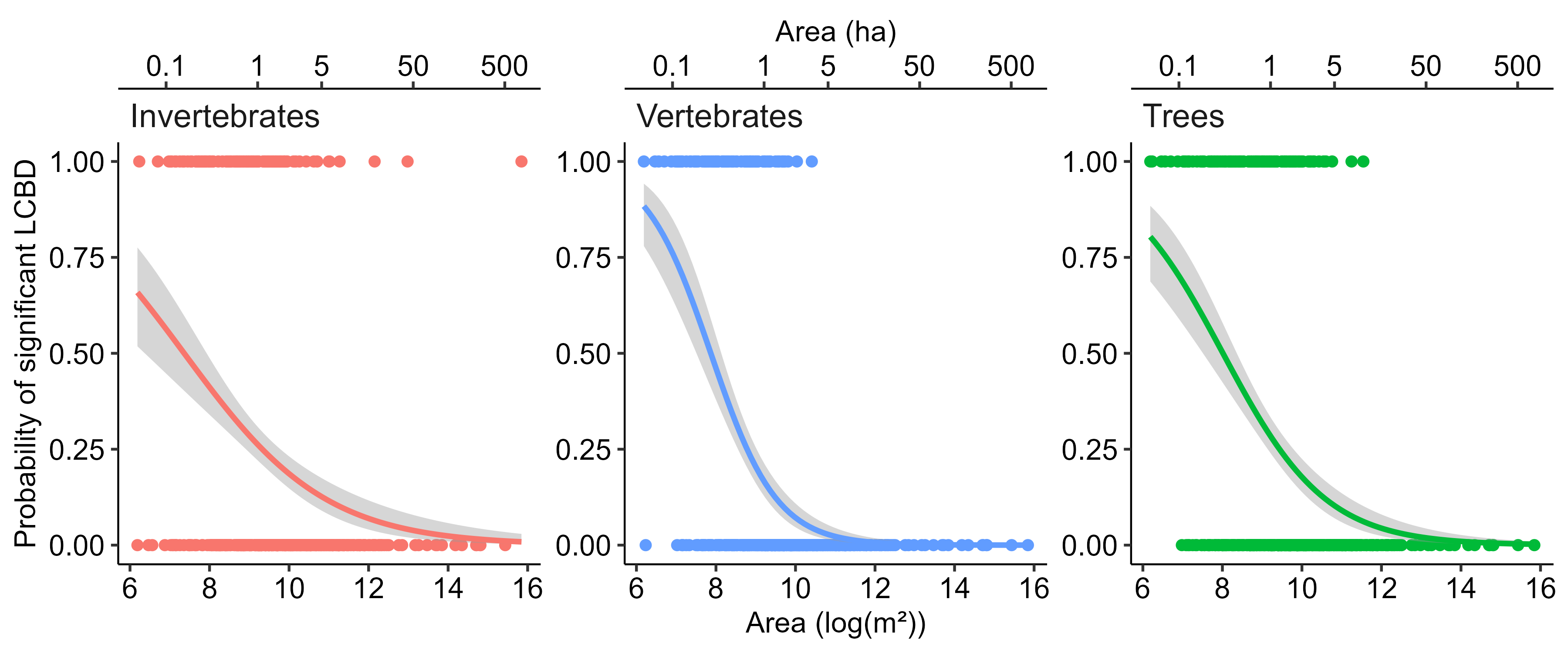
*Figure S9: Logistic regression models showing the probability that patches have significant LCBD values as a function of patch area (log-transformed). Separate curves are shown for invertebrates, vertebrates, and trees (color-coded), with solid lines representing model fits and shaded ribbons indicating standard errors.*

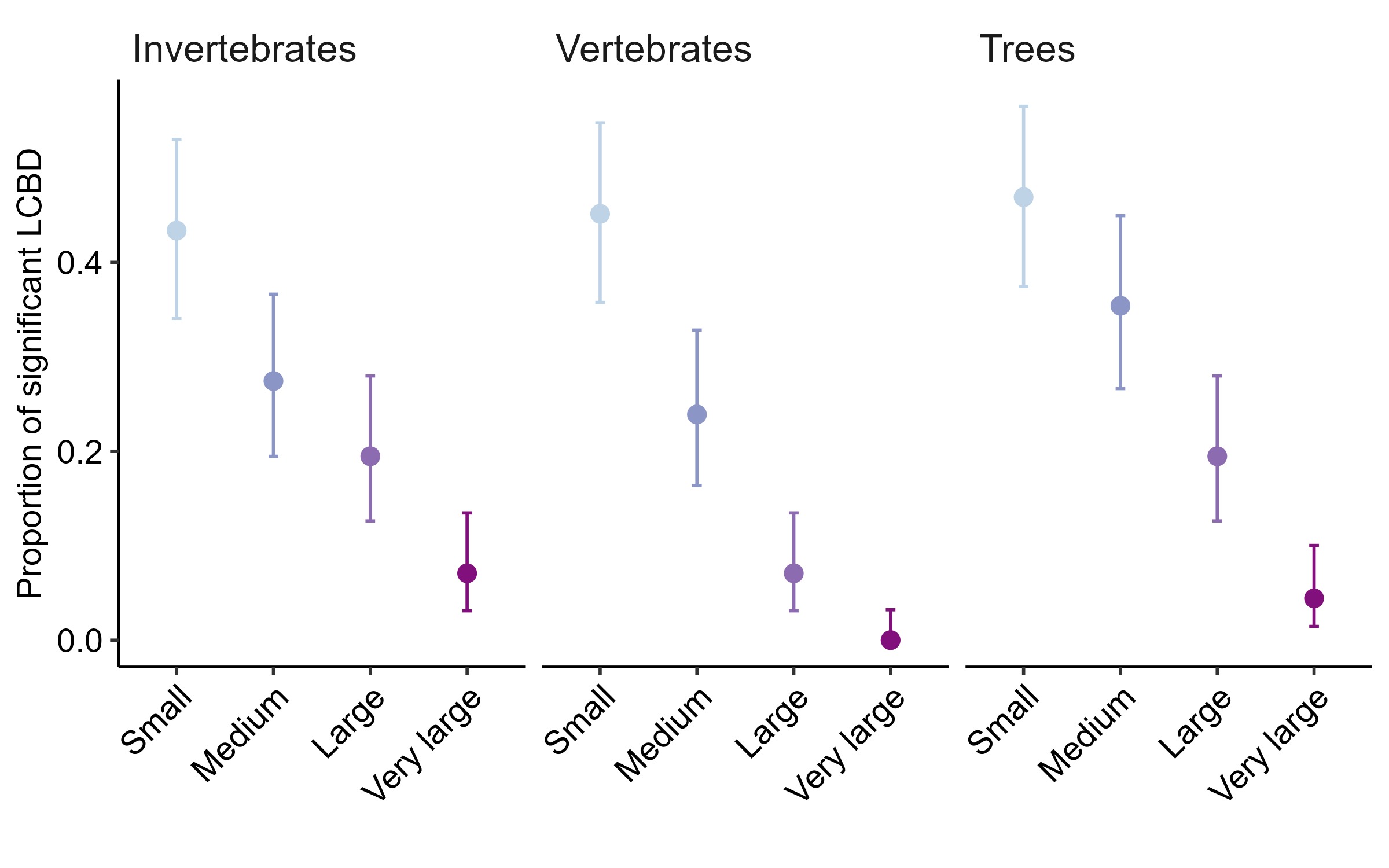
*Figure S10: Proportion of patches with significant LCBD values for each taxonomic group, grouped by patch area quartiles: Small (0–25%), Medium (25–50%), Large (50–75%), and Very Large (75–100%). Dots and error bars represent the mean and standard deviation within each quartiles.*

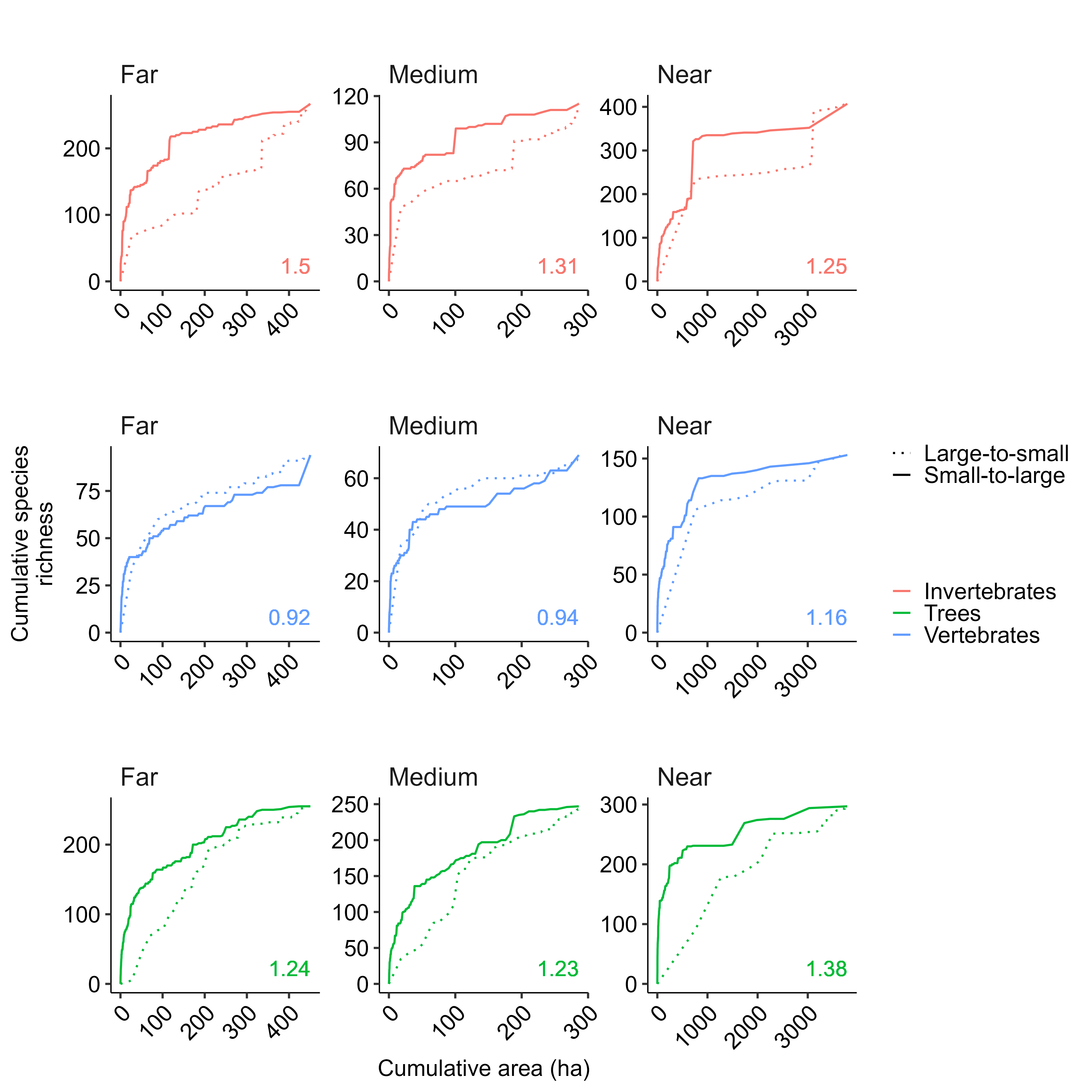
*Figure S11: SLOSS analysis showing cumulative species richness when adding small (dashed) vs. large (solid) patches first for invertebrate, vertebrate, and tree biodiversity (color-coded). Analyses are stratified by patch proximity to forest habitat (Near, Medium, and Far), based on quantiles of distances to the nearest forest patches. SLOSS-index >1 indicates stronger contribution from small patches.*

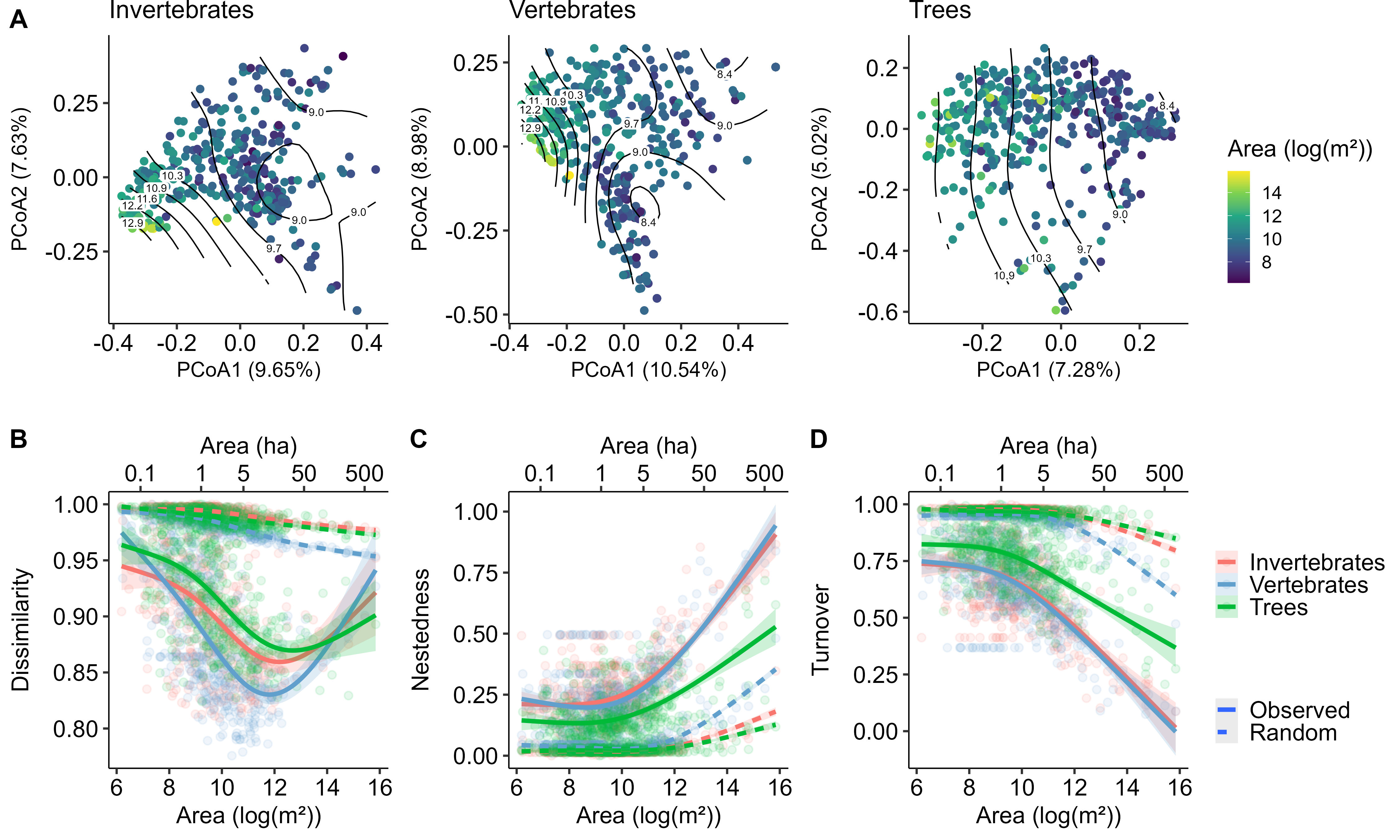
*Figure S12: GAM regressions between log-transformed patch area and multiple alpha and beta diversity metrics across sites, with each dot representing a site and color-coded by taxonomic group. Panels include: (A) log-transformed species richness, (B) average pairwise Jaccard dissimilarity, (C) nestedness component of Jaccard dissimilarity, and (D) turnover component of Jaccard dissimilarity. For panels B–D, dashed lines represent expected values from null models.*

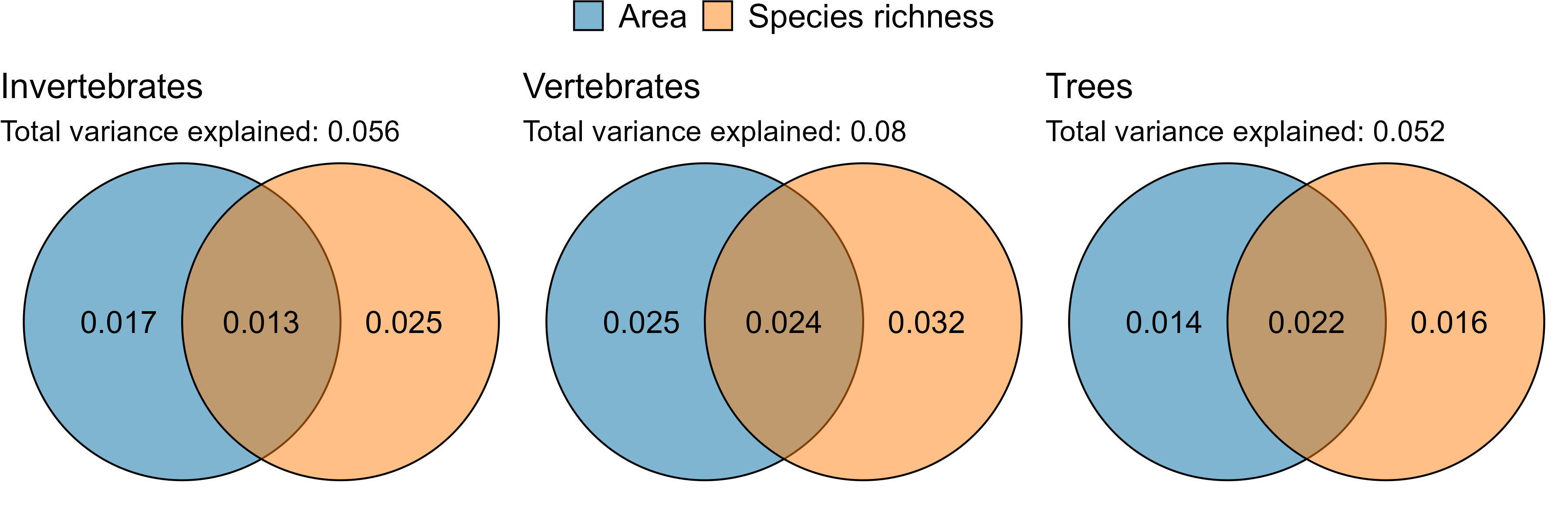
*Figure S13: Venn diagrams of variance partitioning, showing variance explained in Jaccard dissimilarity by area only, species richness only, and their shared contribution (color-coded); labels indicate R^2^ values.*

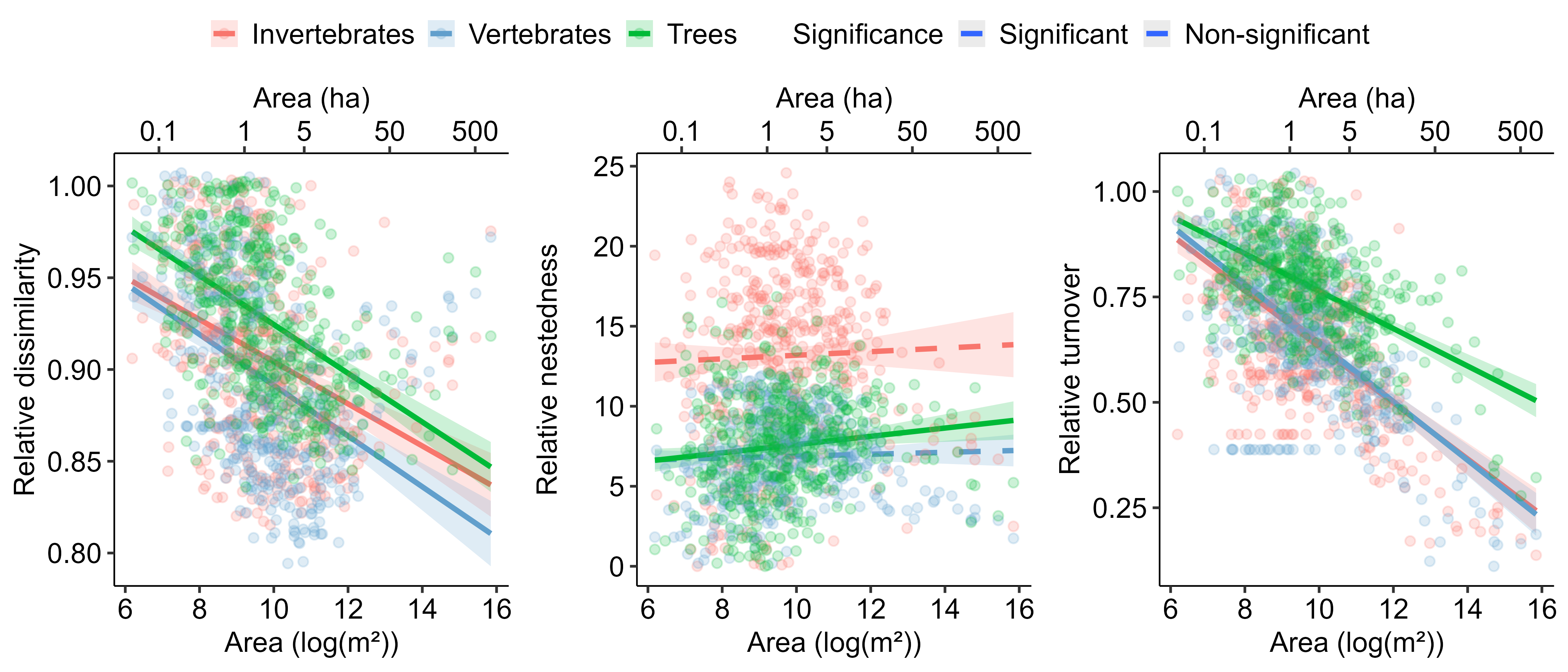
*Figure S14: Regression plots illustrating the relative beta diversity components—Jaccard dissimilarity, nestedness, and turnover—each scaled by their respective null model averages (observed divided by randomized values)—against patch area (log-transformed), for each taxonomic group (color-coded). Each dot represents a site. Solid lines indicate statistically significant linear relationships, while dashed lines denote non-significant ones. Significant relationships suggest that the observed correlation with patch area reflects disproportionate changes in beta diversity components beyond what would be expected from changes in species richness alone.*

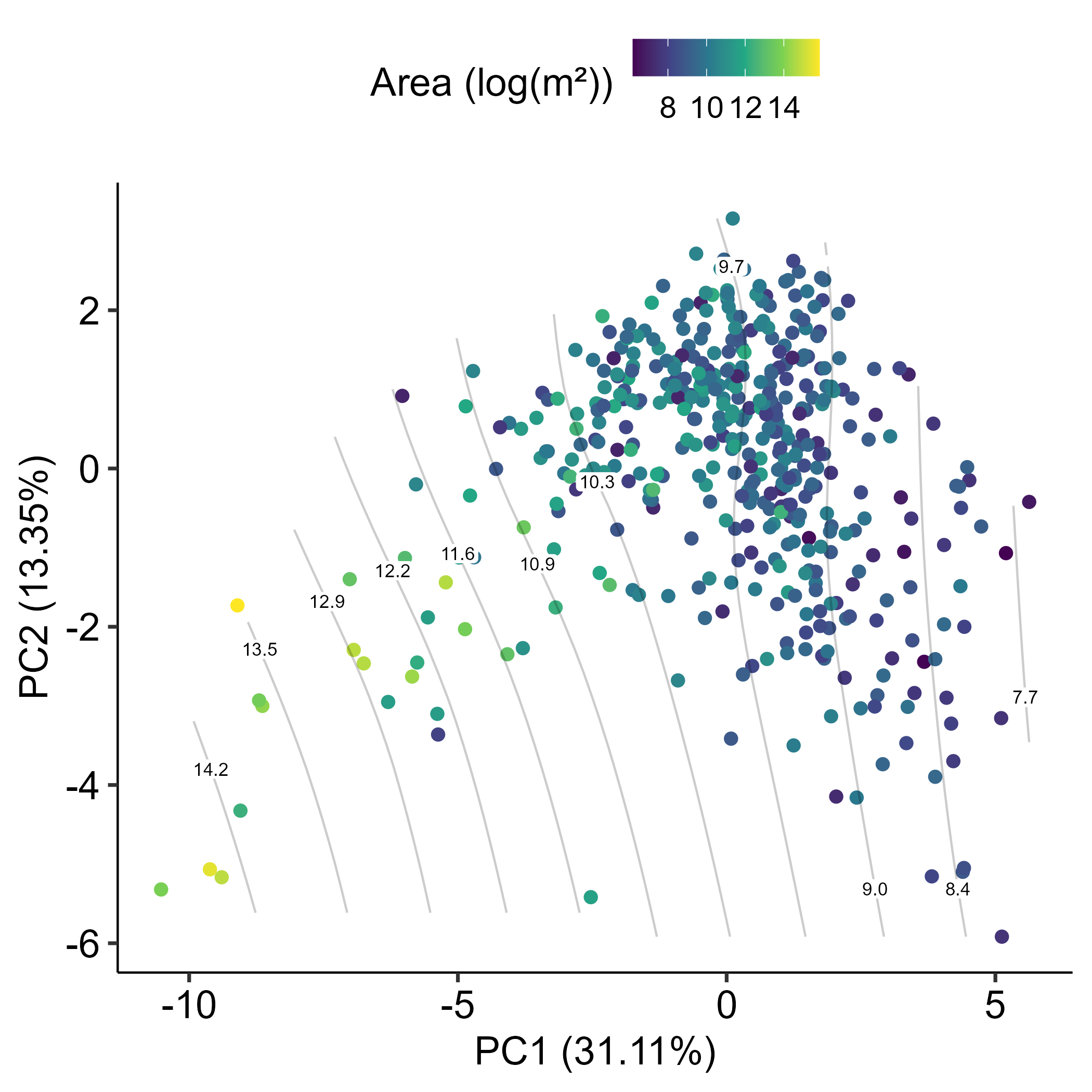
*Figure S15: PCA ordination between environmental factors for each patch and their relation to patch area (log-transformed; color-coded). Smooth surfaces represent fitted GAMs illustrating PCA distances between environmental factors to area (log-transformed).*

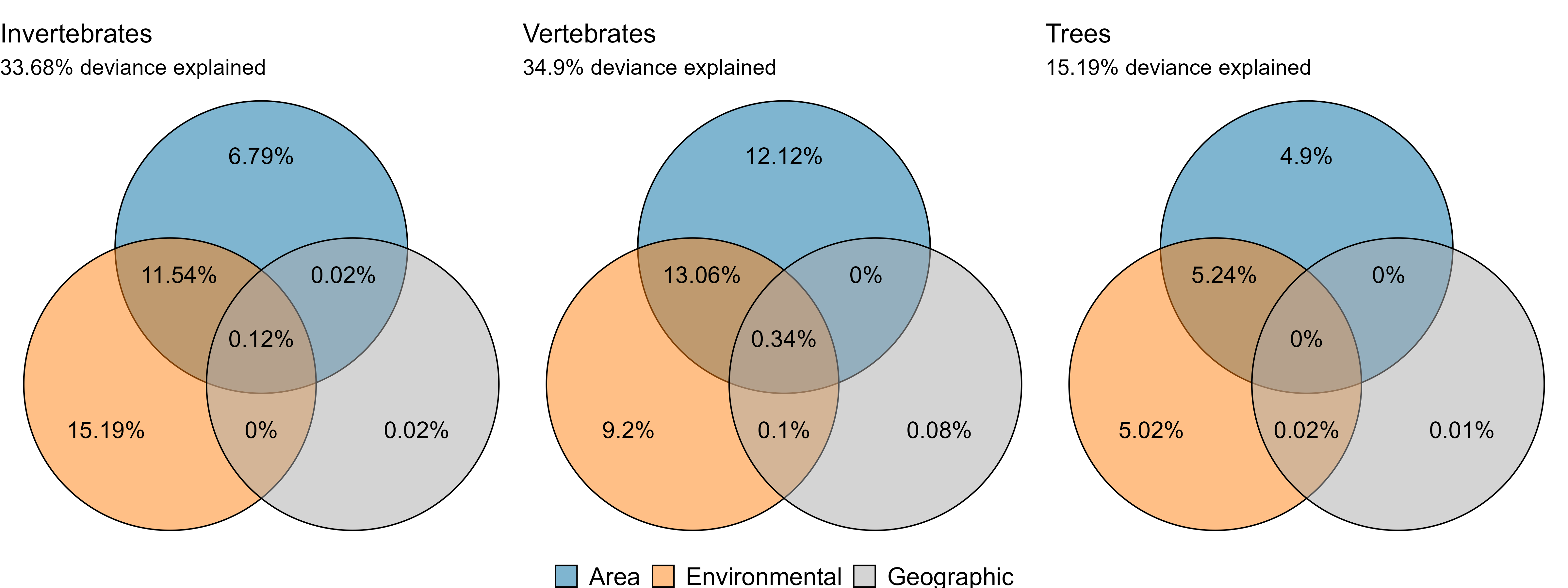
*Figure S16: Venn diagrams depicting the deviance explained by GDM for invertebrates, vertebrates, and trees. The total explained deviance is partitioned into contributions from area, environmental, and geographic factors (color-coded). Percentages indicating the deviance explained by individual factors and their combinations are annotated within the respective sections of the diagrams.*

*Tables S1: Key R packages used in the analyses, including their purpose, version, and relevant references. All scripts and code are available on Zenodo* with the identifier ***.

| *Community analyses* | *vegan v2.6-10*  *adespatial v.0.3-28* | *Oksanen et al. (2025)*  *Dray et al. (2025)* |
| --- | --- | --- |
| *Covariate calculations* | *landscapemetrics v2.2.1*  *exactextractr v0.10.0*  *lidaRtRee v4.0.8* | *Hesselbarth et al. (2019); Baston and ISciences (2023)*  *Monnet (2023)* |
| *Modeling* | *mgcv v1.9-3*  *gdm v1.6.0-7* | *Wood (2025)*  *Fitzpatrick et al. (2025)* |
| *Visualization* | *ggplot2 v3.5.2* | *(Wickham et al. 2025)* |

*Tables S2: Outputs of species richness models as a function of patch area (Area) and isolation from forests (d2_Forest) for each taxonomic group. For each fitted parameter, estimates, standard errors, t-values, and p-values are provided. Significant parameters (p <0.05) are shown in bold.*

|  | Parameter | Estimate | Std. Error | t value | p-value |
| --- | --- | --- | --- | --- | --- |
| Invertebrates | **Intercept** | **-3.71** | **0.34** | **-10.93** | **<0.0001** |
|  | **Area** | **0.55** | **0.03** | **17.14** | **<0.0001** |
|  | d2_Forest | 0.00 | 0.00 | 1.79 | 0.0744 |
| Vertebrates | **Intercept** | **-4.21** | **0.21** | **-20.09** | **<0.0001** |
|  | **Area** | **0.59** | **0.02** | **29.51** | **<0.0001** |
|  | d2_Forest | 0.00 | 0.00 | 0.45 | 0.655 |
| Trees | **Intercept** | **-2.81** | **0.34** | **8.37** | **<0.0001** |
|  | **Area** | **0.49** | **0.03** | **15.18** | **<0.0001** |
|  | **d2_Forest** | **0.00** | **0.00** | **2.63** | **0.00873** |

*Tables S3: Outputs of LCBD models as a function of patch area (Area) and isolation from forests (d2_Forest) for each taxonomic group. For each fitted parameter, estimates, standard errors, t-values, and p-values are provided. Significant parameters (p <0.05) are shown in bold.*

|  | Parameter | Estimate | Std. Error | t value | p-value |
| --- | --- | --- | --- | --- | --- |
| Invertebrates | **Intercept** | **2.79e-03** | **7.91e-05** | **35.23** | **<0.0001** |
|  | **Area** | **-6.11e-05** | **7.55e-06** | **-8.09** | **<0.0001** |
|  | d2_Forest | 4.18e-08 | 2.70e-08 | 1.55 | 0.12 |
| Vertebrates | **Intercept** | **3.08e-03** | **8.21e-05** | **37.51** | **<0.0001** |
|  | **Area** | **-9.00e-05** | **7.84e-06** | **-11.49** | **<0.0001** |
|  | d2_Forest | 1.39e-08 | 2.81e-08 | 0.49 | 0.62 |
| Trees | **Intercept** | **2.93e-03** | **6.42e-05** | **45.54** | **<0.0001** |
|  | **Area** | **-7.40e-05** | **6.13e-06** | **-12.06** | **<0.0001** |
|  | d2_Forest | 9.30e-09 | 2.20e-08 | 0.42 | 0.67 |
